# Linewidth-related bias in modelled concentration estimates from GABA-edited ^1^H-MRS

**DOI:** 10.1101/2024.02.27.582249

**Authors:** Alexander R. Craven, Tiffany K. Bell, Lars Ersland, Ashley D. Harris, Kenneth Hugdahl, Georg Oeltzschner

## Abstract

J-difference-edited MRS is widely used to study GABA in the human brain. Editing for low-concentration target molecules (such as GABA) typically exhibits lower signal-to-noise ratio (SNR) than conventional non-edited MRS, varying with acquisition region, volume and duration. Moreover, spectral lineshape may be influenced by age-, pathology-, or brain-region-specific effects of metabolite T_2_, or by task-related blood-oxygen level dependent (BOLD) changes in functional MRS contexts. Differences in both SNR and lineshape may have systematic effects on concentration estimates derived from spectral modelling.

The present study characterises the impact of lineshape and SNR on GABA+ estimates from different modelling algorithms: FSL-MRS, Gannet, LCModel, Osprey, spant and Tarquin. Publicly available multi-site GABA-edited data (222 healthy subjects from 20 sites; conventional MEGA-PRESS editing; TE = 68 ms) were pre-processed with a standardised pipeline, then filtered to apply controlled levels of Lorentzian and Gaussian linebroadening and SNR reduction.

Increased Lorentzian linewidth was associated with a 2-5% decrease in GABA+ estimates per Hz, observed consistently (albeit to varying degrees) across datasets and most algorithms. Weaker, often opposing effects were observed for Gaussian linebroadening. Variations are likely caused by differing baseline parametrization and lineshape constraints between models. Effects of linewidth on other metabolites (e.g., Glx and tCr) varied, suggesting that a linewidth confound may persist after scaling to an internal reference.

These findings indicate a potentially significant confound for studies where linewidth may differ systematically between groups or experimental conditions, e.g. due to T_2_ differences between brain regions, age, or pathology, or varying T_2_* due to BOLD-related changes. We conclude that linewidth effects need to be rigorously considered during experimental design and data processing, for example by incorporating linewidth into statistical analysis of modelling outcomes or development of appropriate lineshape matching algorithms.

**Highlights:**

- In-vivo GABA-edited ^1^H-MRS data from 222 subjects were filtered to simulate varying linewidth and SNR conditions
- Filtered datasets were quantified with six different modelling algorithms to assess the impact of linewidth and SNR on the metabolite level estimates.
- Synthetic spectra with controlled GABA+ levels and in-vivo-like background signals (applied incrementally) were also assessed.
- For both in-vivo and synthetic datasets, GABA+ estimates showed a significant association with Lorentzian linewidth across most algorithms, even for small changes in linewidth.
- Weaker, often opposing associations were observed for Gaussian linebroadening.
- This indicates a potentially significant confound for studies where linewidth or lineshape may be expected to differ, even slightly, between groups.
- The need for appropriate strategies to account for lineshape differences is highlighted.

Graphical Abstract

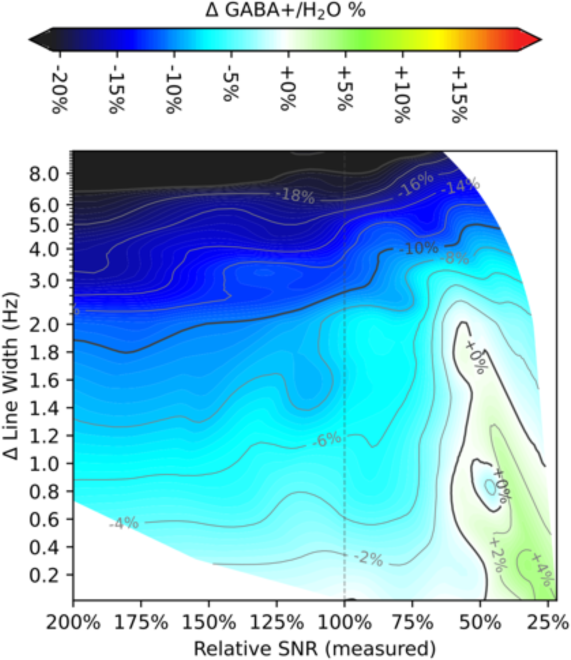

To assess the degree to which aspects of linewidth, lineshape and SNR may confound GABA+ estimates, a collection of in-vivo datasets were quantified with six modelling algorithms, with linebroadening and SNR varied experimentally. Most algorithms showed a strong association between GABA+ estimate and Lorentzian linebroadening (2-5% decrease per Hz), with weaker effects for Gaussian broadening. This indicates a potentially significant confound in cases of differing relaxation parameters between groups or experimental conditions.

## 1 Introduction

The impact of spectral quality on metabolite concentration estimates from MR spectroscopy (MRS) data is a vital yet under-investigated topic, particularly in the context of specialised editing sequences such as MEGA-PRESS^1,2^, frequently used for the assessment of γ-aminobutyric acid (GABA). The linewidth, lineshape^3,4^ and signal-to-noise ratio (SNR) of MRS data can vary substantially depending on acquisition parameters, anatomical region of the voxels, and subject groups. This may impact fitting outcomes and therefore compromise the reproducibility and reliability of metabolite estimation^5–9^. Prior studies have investigated these factors for non-edited PRESS^10^, JPRESS^11^, LASER^12,13^, PEPSI^14^ and STEAM^15,16^ data, and outcomes from a multi-site fitting challenge of synthetic non-edited PRESS data^17^ identified linewidth-related trends in certain conditions. Another recent study^18^ specifically assessed the impact of SNR and linewidth on a synthetic MEGA-PRESS dataset for two of the algorithms considered herein. However, many existing studies report reliability using absolute (unsigned) metrics, reflecting the *magnitude* of estimation errors without regard to direction – perhaps leading to the fallacious assumption that estimation error is normally distributed, without appreciable bias. Although informative, these metrics mask a potentially more severe issue: the risk of *systematic* differences in GABA+ estimation as a function of signal quality, which may bias outcomes and lead to spurious findings if not adequately considered. Many studies also fail to differentiate between Lorentzian and Gaussian broadening mechanisms.

Factors of signal quality become particularly critical in the case of time-resolved or functional MR spectroscopy (fMRS), where metabolic dynamics and design of the functional paradigm may further constrain the number of transients contributing to the resultant spectra, and hence the achievable SNR. Moreover, local variations in effective transverse relaxation rate (T_2_*) associated with blood oxygen level dependent (BOLD) changes are known to have an impact on spectral linewidth. In this context, linewidth-related biases have been demonstrated previously^19,20^, albeit limited to Lorentzian effects on unedited MRS at higher field strengths.

In non-functional contexts, the competing demands of increased anatomical specificity and reduced scan times lead to the question of just how much of a compromise in SNR may be acceptable. Within the same subject, tissue composition and homogeneity, achievable shim quality and hence spectral lineshape will vary between different brain regions. Between subjects or sessions, factors such as age, medication and pathology may also have an impact, as will hardware and sequence parameters between different sites and different study protocols. In all these scenarios, the ability to robustly compare concentration estimates requires rigorous characterisation of any associated biases, and strategies to account for these.

The present study focuses on the impact of linewidth and SNR variation on concentration estimates from GABA-edited MRS data, obtained using the MEGA-PRESS sequence^1,2^. A number of modelling algorithms are available for the processing and quantification of this data, adopting different strategies for separating the signals of interest from overlapping metabolite and background signal. Variability between algorithms has been investigated previously for short-echo-time data ^9,15,21,22^, and for GABA-edited MRS data ^23^. However, for many of these algorithms a comprehensive investigation of the impact of signal quality is currently lacking.

This study therefore aims to characterise the nature and extent of linewidth- and SNR-related effects on concentration estimates obtained from GABA-edited MRS data, and the degree to which these effects vary across sites or according to choice of modelling algorithm. The study compares estimates of GABA+ (GABA modelled together with underlying co-edited macromolecule signals) quantified using six modelling algorithms: FSL-MRS^24^, Gannet^25^, LCModel^26^, Osprey^27^, spant^28^ and Tarquin^29,30^, building on earlier work^23^ comparing several of these algorithms. In-vivo data were filtered to simulate acquisitions of lower SNR (by reducing the number of transients contributing to the averaged spectra for modelling) and increased linewidth (both by Lorentzian and Gaussian linebroadening); results are primarily considered in relation to the unfiltered case. Complementary to the in-vivo findings, a series of synthetic datasets were generated with controlled metabolite linewidth and nominal GABA+ content, progressively incorporating additional confounding factors to independently assess the impact of background/noise signal and co-edited macromolecules on the observed outcomes.

## 2 Methods

### 2.1 In-vivo datasets

The in-vivo datasets underlying the present analysis were obtained from the Big GABA ^31,32^ repository on NITRC, https://www.nitrc.org/projects/biggaba; these datasets have been extensively characterised in previous studies^23,31–33^. Data originated from twenty 3T MRI scanners, each at a different site, covering three major manufacturers (GE, Philips, Siemens). Full details on the hardware and software configurations at each site are described in the previous studies ^23,31–33^, along with details on sample composition and acquisition protocol; an MRSinMRS checklist^34^ and other key details are reproduced in Supplementary Tables 1 and 2.

The datasets include GABA-edited spectra (TR/TE = 2000/68 ms, 320 averages, with editing pulses at 1.9/7.46 ppm for edit-ON/-OFF respectively) acquired from a 3 x 3 x 3 cm^3^ voxel in the posterior cingulate region, along with water-unsuppressed reference data (8 or 16 averages) from the same region. Tissue fractions were derived previously^31^ from T_1_-weighted structural images. The present sample consists of data from 222 adult volunteers in the 18-36 years age range, with approximately even female/male split and having no known neurological or psychiatric illness. Datasets were acquired in accordance with ethical standards of the respective local institutional review boards, with those subjects examined herein having consented to sharing of deidentified data for further study.

#### 2.1.1 In-vivo data preparation

After initially loading with import functions and coil combination from Gannet^25^, in-vivo spectral data were processed using a common pipeline built on FID-A^35^ functionality and described fully in an earlier study^23^. The basic pipeline includes rejection of motion-corrupted transients, alignment of individual transients by spectral registration^36^, averaging, eddy-current correction^37^, zero-order phase adjustment of each sub-spectrum according to a dual-Lorentzian model for creatine (Cr) and choline (Cho), before final alignment between edit-ON and edit-OFF sub-spectra, again using spectral registration^36^.

Datasets were subsequently filtered to simulate spectra of varying quality. After initial loading, metabolite data were sub-sampled to reduce the effective number of averages, and thereby the SNR. Random sampling of transients was performed after spectral registration, immediately before averaging, retaining adjacent edit-ON/edit-OFF pairs but without attention to phase cycling. Randomly sampled subsets proportional to [all, 1/2, 1/4, 1/8, 1/16, 1/32] of the available data (i.e., [320, 160, 80, 40, 20, 10] averages) were selected.

To simulate acquisitions of varying linewidth, Lorentzian linebroadening (LB_lorentz_) was applied to the resultant spectra from each of the randomized subsets, by multiplication of the time-domain FID by an exponential decay function 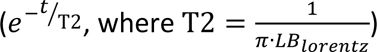. LB was applied at 0-2 Hz in steps of 0.2 Hz, 2-5 Hz in steps of 0.5 Hz, and 5-10 Hz in steps of 1 Hz. A similar approach was used to separately assess Gaussian line-broadening (LB_gauss_), multiplying the FID by 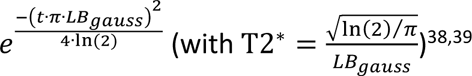. Filtered data were saved at the same resolution as the input data, without zero-fill. No filtering was performed on water reference data.

#### 2.1.2 Quality Control

Quality of the incoming in-vivo data and initial processing was evaluated using only the unfiltered spectra – that is, spectra processed using the full set of available transients (after rejecting motion-corrupted transients) and without additional linebroadening, before fitting. Two rejection criteria (adopted from an earlier study^23^) were applied on a per-dataset basis, designated **R1** and **R2** in subsequent usage; acquisitions which failed either of these criteria were excluded from further analysis:

- **R1** captures spectra having strongly aberrant features in the fit range: processing was deemed to have failed if the 0-lag cross-correlation of the normalized, reconstructed frequency domain difference spectrum in the metabolite range (2.6-4.2 ppm) with the normalized mean of all other difference spectra was below 0.5 or differed from the group mean by more than three standard deviations.
- **R2** establishes thresholds on basic signal quality metrics: linewidth (FWHM > 10 Hz^40^, measured from the negative n-acetyl aspartate (NAA) peak in the difference spectrum (NAA_diff_)), and SNR (< 80, defined by maximum negative peak amplitude of NAA_diff_ in the [1.8, 2.2] ppm interval, over standard deviation across the [-2, 0] ppm range.

The modelling algorithms tested vary greatly in their reported quality-of-fit metrics (%SD, CRLB, FitError, etc); in the context of experimentally manipulated SNR and linewidth, these may present different biases^41^ and limitations. Since our analysis involves deliberately degrading SNR and linewidth (in some cases beyond the range which would ordinarily be considered acceptable), no further rejection criteria were applied to the filtered spectra.

Extreme outliers in the modelled GABA+/H_2_O estimates (see section 2.4) were removed according to a third criterion:

- **R3** rejects extreme outliers from the modelled GABA+/H_2_O estimates. Individual estimates diverging from the median by more than five times the median absolute deviation (MAD) for that algorithm were rejected from subsequent analysis, with median/MAD parameters evaluated on a “higher quality” subset: estimates modelled on the full set of transients with 0-6 Hz linebroadening.

For the simulated datasets (section 2.3), only **R3** was applied, separately for each of the nominal simulated GABA+ concentrations. Outlier-robust methods were also adopted in subsequent analyses, wherever feasible.

### 2.2 Basis Set Preparation

All the assessed fitting algorithms (except Gannet) require prior knowledge in the form of a basis set. For each algorithm, the same vendor-specific basis sets were used, in a format appropriate to the tool. These had been previously created^23,27^ using fast spatially resolved 2D density-matrix simulations^42^ implemented in FID-A^35^, with ideal excitation pulses and vendor-specific refocusing pulses and timings, and using chemical shifts and J-coupling coefficients from Kaiser et al.^43^. Basis sets included components for GABA, Glutamate (Glu), Glutamine (Gln), glutathione (GSH), NAA, n-acetylaspartylglutamate (NAAG), and parametrized Gaussian components representing co-edited macromolecules around 0.91 ppm (denoted MM09ex; FWHM=10.9 Hz), and co-edited macromolecule signal underlying the GABA peak at 3.0 ppm (denoted MM3co) with FWHM = 14 Hz and scaling equivalent to two protons^44^. The sum of GABA and MM3co fit components is reported herein as GABA+.

### 2.3 Synthetic datasets

To independently assess the contribution of overlapping metabolite signal, co-edited macromolecule signal and background signals (baseline, noise, artefactual components) to any observed effects, a series of hybrid spectra were synthesized, combining simulated metabolite components with realistic background signal derived as follows from the in-vivo data. For each of the unfiltered in-vivo spectra, the modelled baseline and fit residuals from each of the basic modelling algorithms in an earlier analysis^23^ were extracted. For each point across the fit range (0.2 - 4.2 ppm) of the frequency domain spectrum, the median baseline and median fit residual across algorithms was evaluated to yield a “consensus” background signal for each dataset. This signal represents features of the in-vivo data which were consistently *not* covered by the prior knowledge metabolite models of the algorithms investigated: broad baseline fluctuations, co-edited macromolecule or metabolite contributions not captured by the model, artefactual components, and random noise. By considering background derived by multiple different fitting models, we reduce the risk that the background signal thereby derived, in combination with simulated basis spectra, is perfectly conformant to any one of the algorithms assessed^45^. The across-algorithms median frequency domain baseline model signal will hereafter be referred to as the “consensus baseline”, describing broad undulations in the in-vivo spectrum, while the across-algorithms median fit residual will be referred to as the “consensus residual”, describing signals in the fit range not captured by existing metabolite and baseline models. The combination of these will be referred to as the “consensus background”. These parts are illustrated for a representative subject in Figure 1 (A).

**Figure 1.**
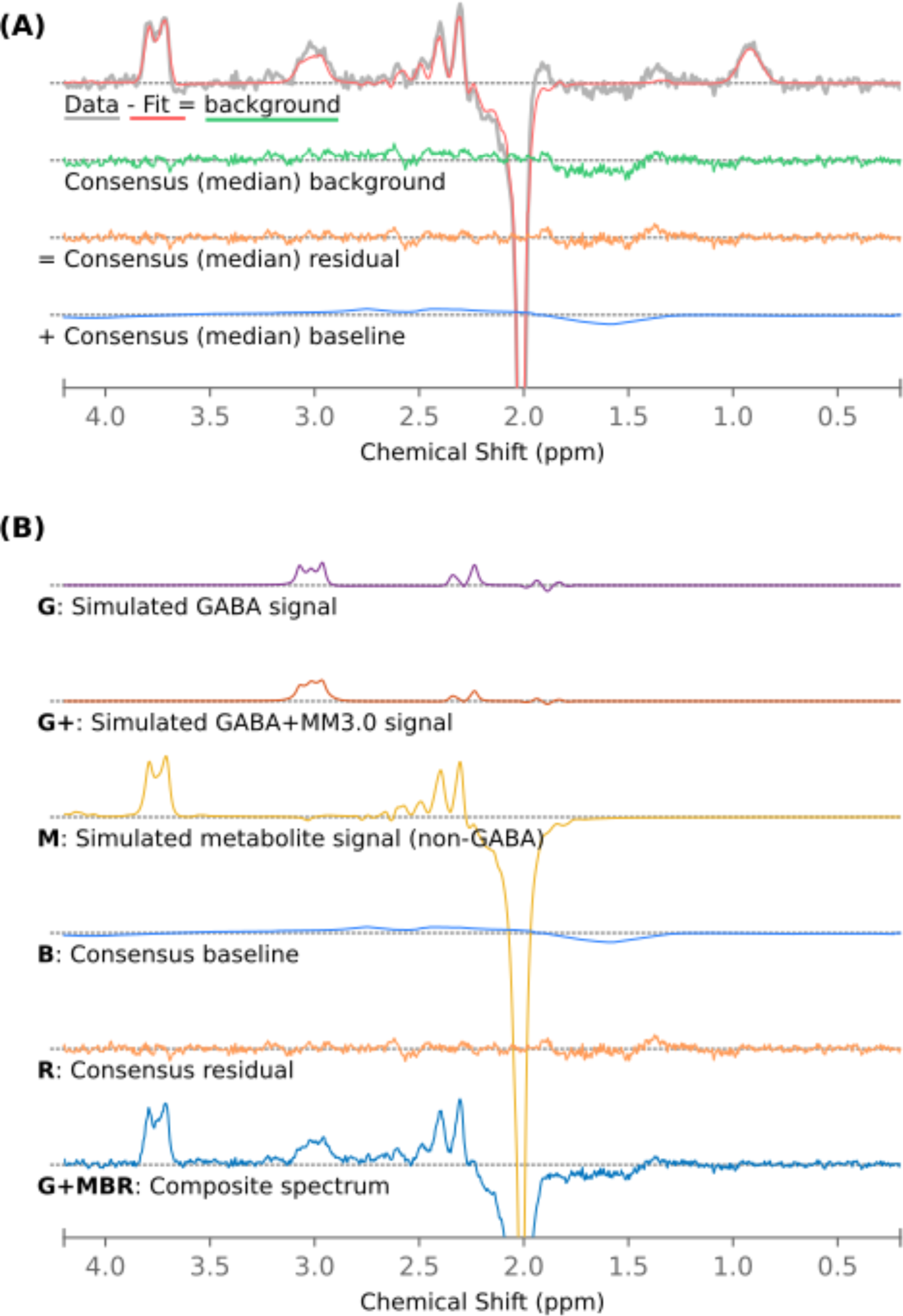
(A) Derivation of consensus background, residual and baseline signal: median across algorithms from a single representative in-vivo subject, subsequently combined with simulated metabolite components (B) to yield in-vivo-like simulated datasets.

Background signals from thirty subjects were selected for further analysis: ten typical, representative backgrounds from each of the three hardware manufacturers, randomly sampled from the upper 70-percentile correlation of the subjects’ consensus background signal with the group mean for the corresponding manufacturer. For each of these thirty background signals, simulated metabolite signal components were added from the corresponding manufacturer-specific basis set. Signal components were combined incrementally in the following schemes, illustrated in Figure 1 (B):

- **G**: **G**ABA only (no background), in simulated concentrations of [2.5, 3.2, 3.9] mM.
- **GM**: **G**ABA as above, with a standard set of simulated **M**etabolites: 12.5 mM NAA, 2.5 mM NAAG, 6 mM Cr, 4 mM phosphocreatine (PCr), 1.2 mM glycerophosphorylcholine (GPC), 1.8 mM phosphorylcholine (PCh), 7.5 mM *myo*-inositol (mI), 5 mM lactate (Lac), 9 mM Glu, 3.5 mM Gln, 1 mM GSH.
- **G+M**: **G**ABA**+**, simulated as GABA plus a fixed 1.5 mM MM3co, underlying 3.0 ppm macromolecule signal, in *total* concentrations of [2.5, 3.2, 3.9] mM, with a standard set of **M**etabolites as above.
- **GMB**: **G**ABA with a standard set of **M**etabolites (per GM case), with consensus **B**aseline signal included.
- **GMR**: **G**ABA with a standard set of **M**etabolites (per GM case), with consensus **R**esidual signal included.
- **GMBR**: **G**ABA with a standard set of **M**etabolites (per GM case), with both consensus **B**aseline and consensus **R**esidual signals included.
- **G+MBR**: A complete in-vivo-like signal: **G**ABA**+** (GABA with a fixed 1.5 mM MM3co), in total concentrations of [2.5, 3.2, 3.9] mM, with standard set of **M**etabolites as above; consensus **B**aseline and consensus **R**esidual signals included.

For each of these cases, metabolite linewidth was varied from 2 - 12 Hz in steps of 0.5 Hz (Lorentzian linebroadening); background and reference signals were not affected. Hence: 21 linewidths for each of three GABA(+) concentrations were quantified using seven different combinations of metabolite and background: a total of 441 simulated spectra for each of the thirty selected subject backgrounds.

### 2.4 Processing and Quantification

Processed and filtered spectra (both in-vivo and simulated) were passed into each algorithm in an appropriate format. In general, the algorithms were invoked using the developer-supplied default or recommended configuration parameters for GABA-edited MEGA-PRESS data, as detailed in a prior study^23^ and summarised in Supplementary Section H, with basis sets as detailed in section 2.2. Some adjustments to baseline modelling were made for Osprey and LCModel (0.6 ppm spline knot spacing adopted for each), and for spant (see Supplementary Section E, Supplementary Table 7 and Supplementary Figure 10). Built-in processing steps for the various implementations are bypassed; this is particularly significant for Gannet, which would ordinarily perform linebroadening and zero-fill before modelling. Batch processing was automated in Matlab (v2021a, MathWorks Inc., Natick, MA, USA), except for spant which was scripted externally in R ^46^ (v3.5.2). This processing and modelling workflow is summarised in Figure 2.

**Figure 2:**
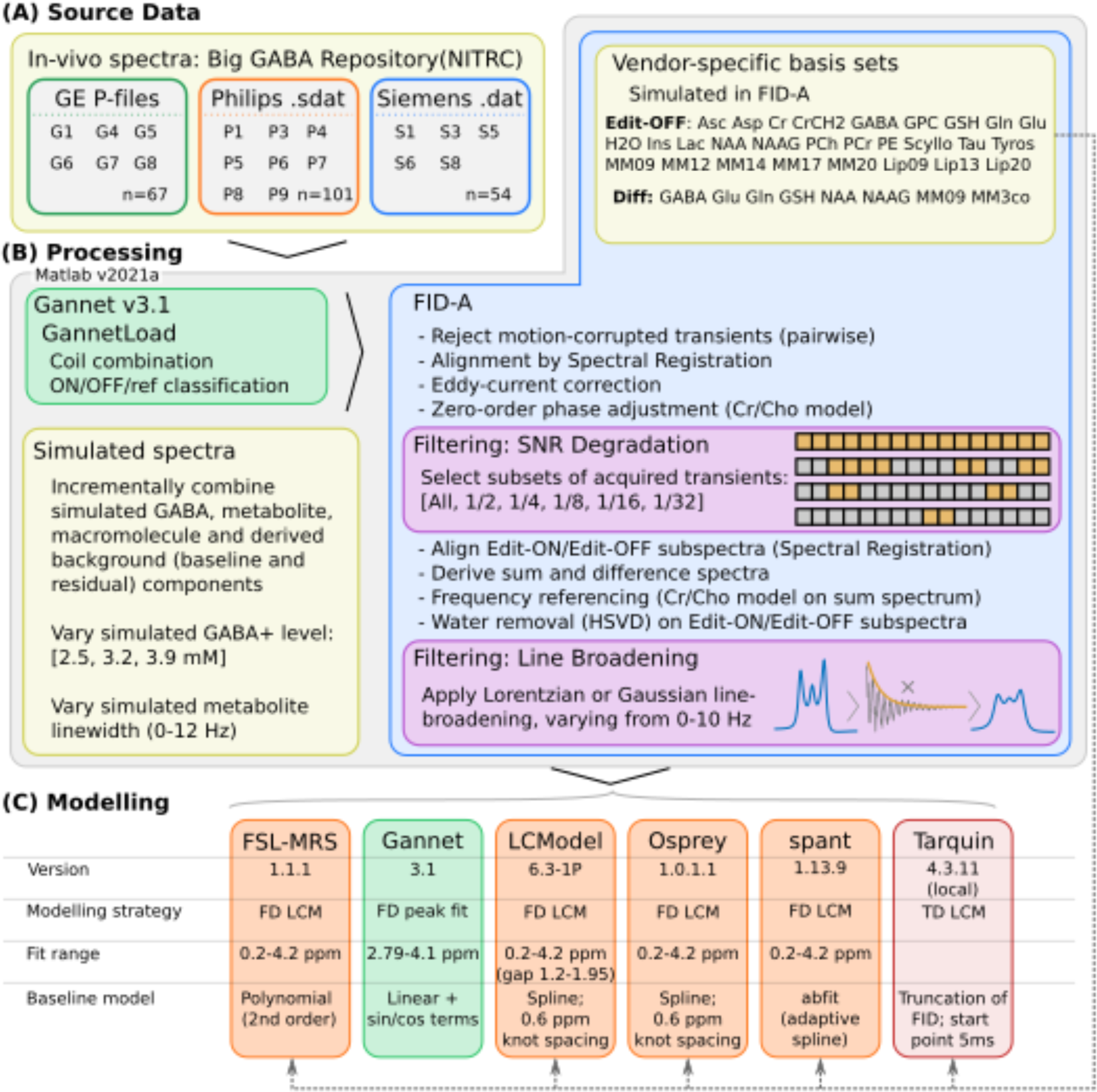
Data (A), processing (B) and modelling (C) workflow, summarising applied filtering and configuration of the various algorithms assessed. Figure derived from Craven et al, 2022^23^.

The raw ratio of metabolite to water signal intensities was scaled with tissue correction^47^ using previously derived tissue fractions^31^; full details on this re-scaling have been documented previously^23^. Tissue corrected molar concentrations scaled to the unfiltered water reference are denoted “/H_2_O”. While the bulk of our analysis focuses on GABA+ from the edited difference spectra, filtered edit-OFF sub-spectra are also modelled for the in-vivo datasets, to allow the efficacy of referencing to total creatine from the edit-OFF sub-spectrum (tCr_off_) to be discussed; associated modelling parameters are supplied in the Supplementary Material, section H.2.

### 2.5 Numerical and Statistical Analysis of Modelling Outcomes

Outcomes of the batch modelling were collated and analysed using locally developed scripts written in Python (v3.9.2), leveraging the pandas ^48^ (v1.5.2) data analysis framework, with numeric methods from NumPy ^49^ (v1.23.5) and statistical methods from the SciPy ^50^ (v1.9.3), pingouin ^51^ (v0.5.2) and statsmodels ^52^ (v0.13.5) libraries. Visualisation was performed using matplotlib ^53^ (v3.3.4) and seaborn ^54^ (v0.12.1).

Where appropriate, normality was assessed with the Shapiro-Wilk method ^55^, and comparability of variance using Fligner-Killeen’s test ^56^; t-tests were performed using Welch’s method^57^. In sub-analyses where Holm-Bonferroni ^58,59^ correction was applied, adjusted p-values are denoted p_holm_, with a corrected significance threshold defined as p_holm_<0.05; uncorrected p-values are denoted p_unc._.

The dataset described herein has previously been assessed with respect to demographics, underlying signal quality and with respect to broad differences with certain combinations of vendor and algorithm ^23,31^. To harmonise estimates across sites (and hence, across vendors), the ComBat approach^60^ was used; individual harmonisation factors were determined from the unfiltered spectra, then applied to the full set of filtered spectra (prior to rejection of outlier estimates, R3). For subsequent analyses, the impact of inter-individual variation was mitigated by expressing in-vivo estimates for each subject and algorithm as percentage differences relative to the estimate from the corresponding unfiltered spectra. For each filtered spectrum, achieved SNR and linewidth were measured using the negative NAA_diff_ peak (as described in section 2.1.2), expressed both as absolute values and as changes relative to the unfiltered spectrum.

Variance Partition Coefficients (VPCs) were derived for subject, SNR and linewidth factors on filtered spectra at various applied SNR and LB levels, using an unconditional linear mixed-effects model implemented in R^46^ with the lme4^61^ package. Within each group (site), a series of paired t-tests was performed, evaluating estimates at each combination of SNR and LB against those obtained from the unfiltered data (*uncorrected*, significance threshold p=0.05); this allowed an indicative rate of spurious findings (type I errors) associated with a particular change in linewidth to be identified.

One limitation of applied line-broadening (essentially a smoothing function) is that this will also alter the SNR of the data; in the present study, numerical methods were used to separate these factors. For each algorithm, a two-dimensional mesh representation of obtained GABA+ values as a function of both the nominal, applied LB and the achieved (measured) SNR was generated. Estimates from each algorithm were distributed evenly into 14 bins according to LB, then within each of these into seven even bins according to relative SNR. Mesh coordinates were defined as the median of SNR and linewidth parameters within each bin. Uniform mesh refinement with cubic interpolation allowed the expected GABA+ estimate at arbitrary combinations of SNR and LW within the experimental range to be approximated – hence, the impact of linewidth to be assessed independently of SNR. The same mesh was used to generate contour maps describing relative change in concentration estimates (presented in Figure 4 and Figure 5), for visual inspection to guide subsequent analysis steps.

The effect of small changes in linewidth on GABA+ estimates, as a function of starting linewidth, was quantified with a piecewise linear description. This was derived by taking the linear least-squares regression of GABA+ on linewidth, as a function of linewidth 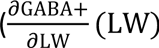, with LW measured after filtering) over a five-point sliding window across the range of linewidths for each individual subject, for each algorithm assessed. The median of 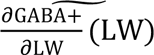 is calculated across a limited range of linewidths: the first 2.12 Hz of broadening for in-vivo data, and nominal linewidth of 5.26 ± 1.06 Hz for the simulated data (guided by the linewidths observed in unfiltered in-vivo data). This factor, which quantifies the approximated linear relation between measured concentration estimate and linewidth (valid over a limited range of linewidths), will hereafter by referred to as the linear linewidth factor, LLWF. This factor forms the basis for statistical comparisons across modelling algorithms and vendors (see section 3.1.2). While the overarching focus of the present analysis remains on GABA+, similar factors were assessed for Glx and tNAA (both from the edited difference spectrum) and tCr*_off_* (from the edit-OFF sub-spectrum).

#### 2.5.1 Synthetic Data

For each of the incremental synthetic models, for each algorithm and at each simulated linewidth, Standard Deviation (SD), Mean Signed Difference (MSD) and Mean Absolute Error (MAE) of GABA+ estimates were evaluated, relative to the nominal (simulated) concentration. The Pearson correlation between nominal and estimated concentration was also assessed.

## 3 Results

### 3.1 In-vivo data

Four datasets were rejected according to criterion **R1**; no further rejections according to criterion **R2**. The global mean estimate for GABA+/H_2_O across all algorithms and subjects was found to be 3.2 ± 0.8 i.u.. Rejection rate according to **R3** is summarised in Supplementary Figures 2A and 3A. Linewidth of the incoming data (after pre-processing, without filtering) was measured as [4.25, 5.33, 7.61] Hz for the [5, 50, 95]-percentiles; per-site SNR and linewidth are presented in Supplementary Table 3.

#### 3.1.1 Fit Outcomes

Mean filtered data and fits by each algorithm for low and high linewidth and SNR cases are presented in Figure 3 and Supplementary Figure 1 (the latter covering the full fit range). Particularly of note here is that the right side of the GABA peak is clearly elevated in broader linewidth cases, as it increasingly merges with broadened Glx signals in the 2.3-2.6 ppm range; for several algorithms, the baseline model is seen to distort upwards in this area.

**Figure 3.**
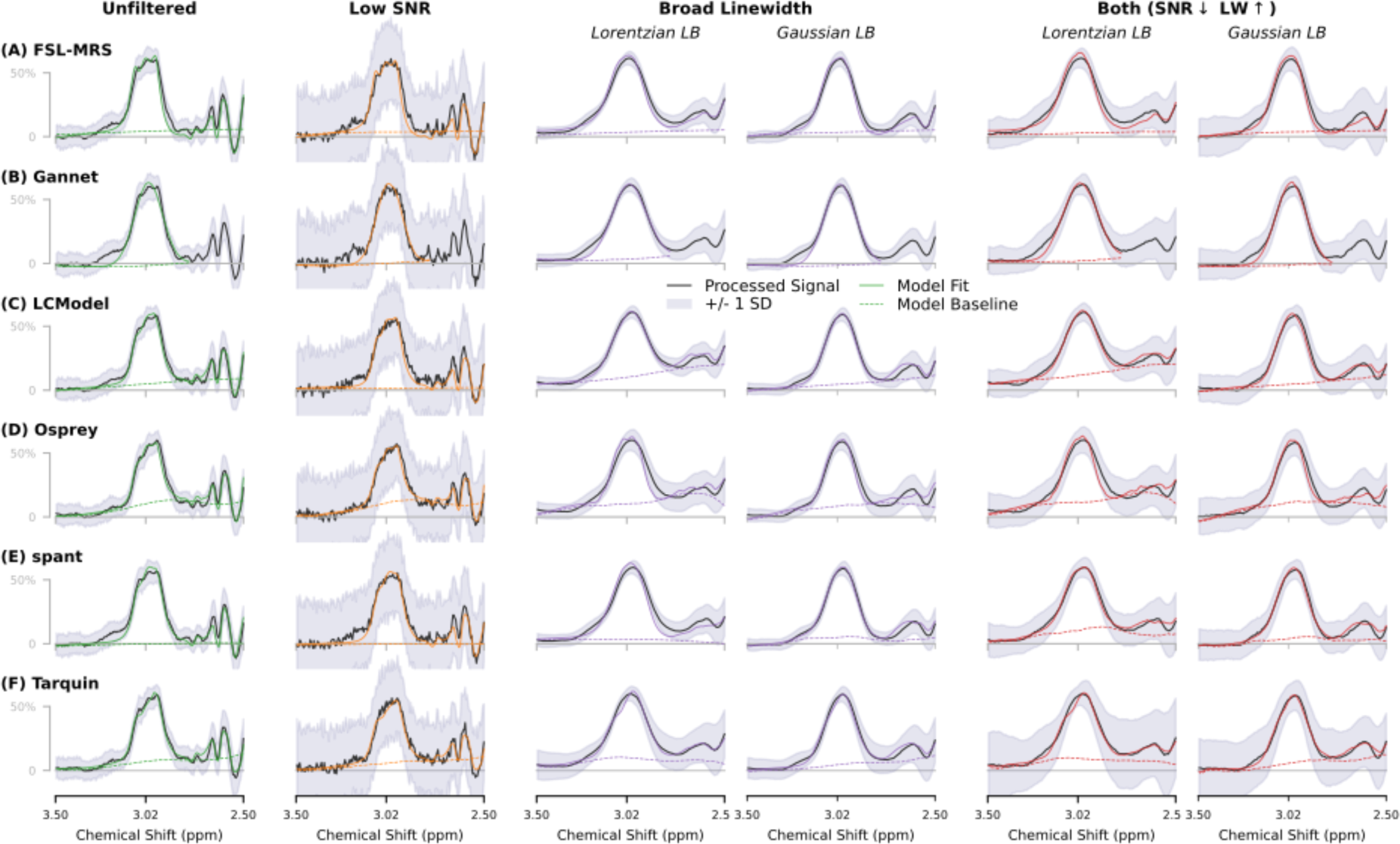
Mean spectra, GABA+ fits and modelled baseline for each algorithm, illustrating the unfiltered original data alongside filtered variants having lower SNR (1/32 subset), higher linewidth (10 Hz of Lorentzian and Gaussian broadening, separately), and a combination of SNR and LB factors. Vertical scaling is normalised. Outcomes over the full fit range are presented in Supplementary Figure 1.

Baseline estimates after Gaussian LB are less variable than after Lorentzian LB case for several algorithms. Also of interest is behaviour around a characteristic hump at 3.2 ppm (not directly attributable to GABA+); most algorithms manage to reject this in the lower linewidth cases (even at lower SNR), but appear less successful at doing so with broader linewidths – potentially dragging the GABA model slightly to the left in the process; LCModel in particular appears to have enveloped essentially all the 3.2 ppm signal. The observed baseline behaviour prompted further exploratory analysis on LCModel baseline knot spacing, described in Supplementary Section G, Supplementary Figure 14 and Supplementary Table 9.

VPCs for subject, LB and SNR factors are presented in Supplementary Table 4; applied Lorentzian LB accounted for 12.8 ± 10.8 % of total variance in GABA+ estimates, depending on algorithm. Applied Gaussian LB accounted for substantially less variance, 2.8 ± 1.6 %, as did the applied SNR factor: 1.0 ± 1.0 %.

Paired t-tests on the obtained GABA+/H_2_O estimates for each site, comparing each SNR/LB step against the *same* data in the unfiltered case, allowed assessing the likelihood of spurious (false positive) findings given certain SNR/LW differences. These outcomes are presented in Supplementary Figure 2C and Supplementary Figure 3C. Several algorithms (LCModel, spant, Tarquin) yielded statistically significant differences (p_unc_ < 0.05) between filtered and unfiltered cases in over half the sites assessed after less than 0.5 Hz of LB_lorentz_; other algorithms did so at a lower rate, with FSL-MRS and Osprey appearing somewhat more robust (but still far from ideal) in this regard, reporting < 20% statistically significant differences within the first 1 Hz of linewidth difference.

#### 3.1.2 LLWF: GABA+ estimate as a function of Linewidth

GABA+ estimate (relative to the unfiltered case) as a function of relative SNR and linewidth is presented in Figure 4 and Figure 5 for Lorentzian and Gaussian LB respectively. All of the algorithms except FSL-MRS show a clear trend towards lower GABA+ estimates with increasing Lorentzian linewidth; visual inspection suggests GABA+ estimates decrease by roughly 10% with 3 Hz of linebroadening for all algorithms except FSL-MRS. The SNR factor does not present a clear trend, although several algorithms (FSL-MRS, Gannet, LCModel, spant) yield slightly higher estimates for the lowest SNR cases. Relative estimates for co-edited Glx are presented for interest in Supplementary Figures 4 and 5, but are not analysed further.

**Figure 4.**
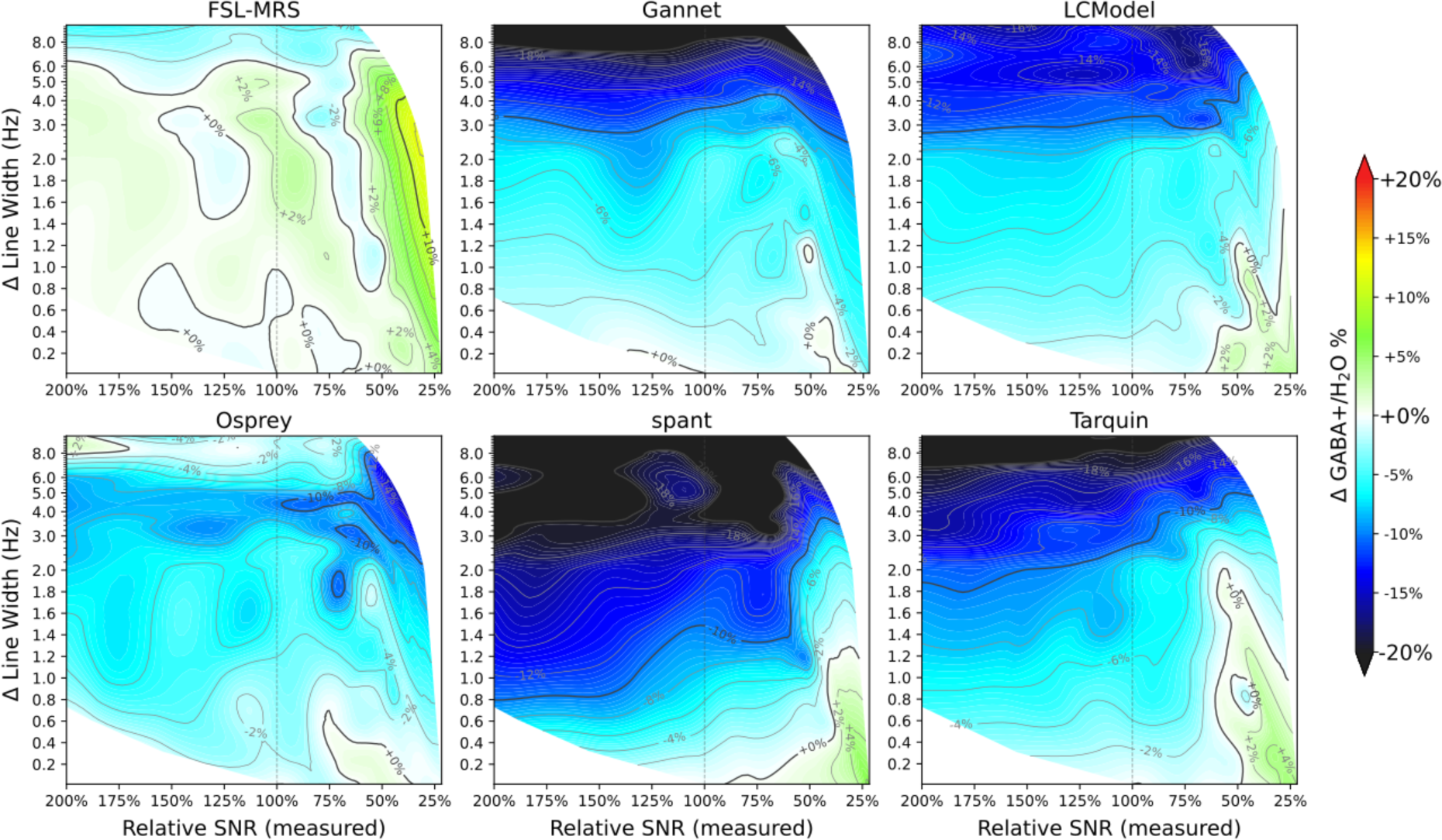
GABA+ estimate (relative to unfiltered case) as a function of Lorentzian linewidth and SNR, for each of the algorithms assessed.

**Figure 5.**
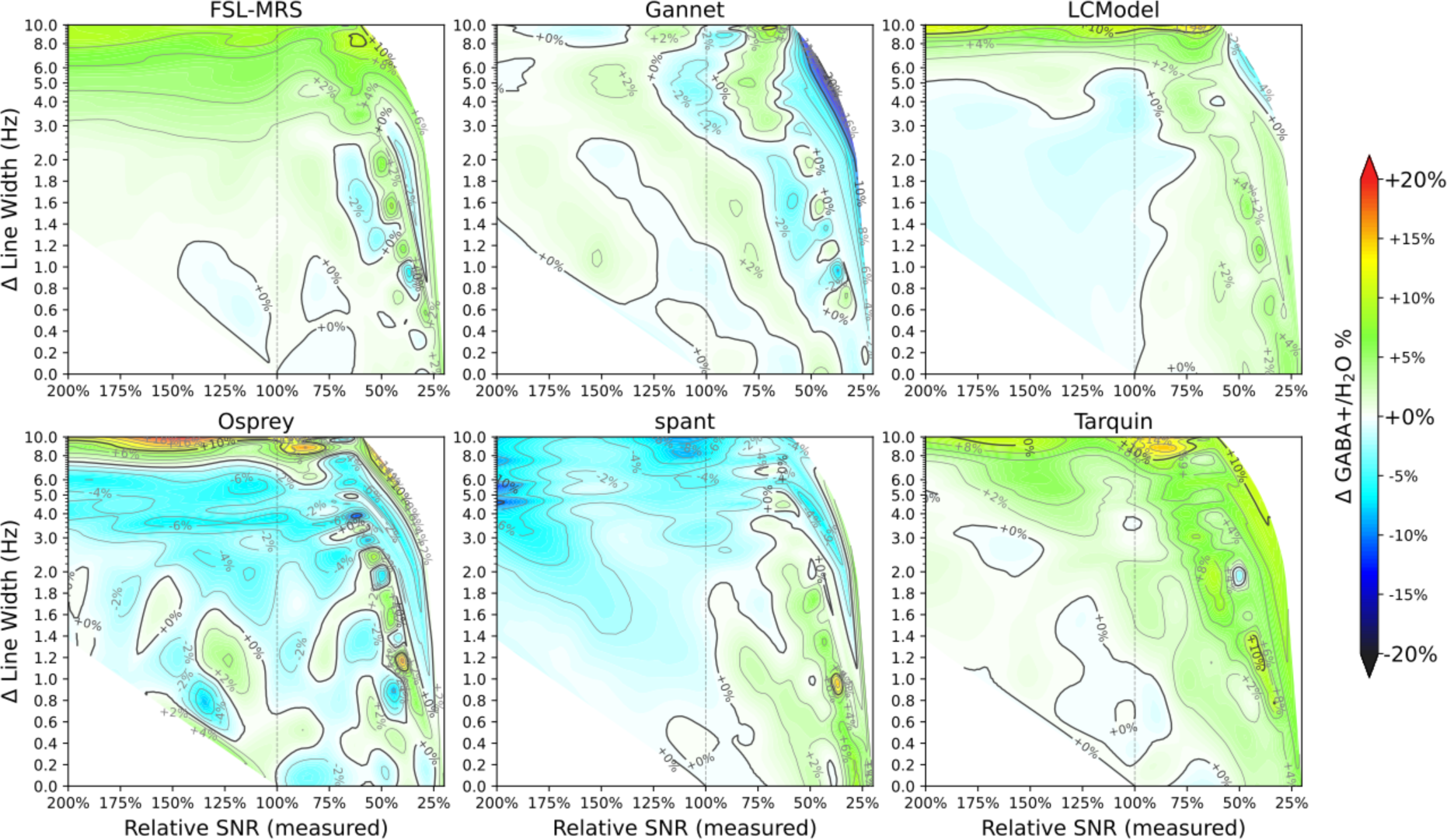
GABA+ estimate (relative to unfiltered case) as a function of Gaussian linewidth and SNR, for each of the algorithms assessed.

Conversely, Gaussian linebroadening shows a more moderate effect on GABA+ estimates, often opposing the Lorentzian effects: there were slight positive associations for moderate LB_gauss_ factors (<1% per Hz LB, for LB_gauss_<4 Hz), becoming more substantial for larger broadening factors). SNR effects are also more apparent in the higher LB_gauss_ cases.

The linewidth relation is explored further in Figure 6. Estimates modelled at a fixed SNR (i.e., with LB and SNR factors separated as described in section 2.5) showed the same trends across all algorithms. These estimates are also presented in Figure 6A, having been sampled across the fixed SNR values (relative SNR 100%, indicated by a dashed vertical line in Figure 4 and Figure 5). Changes in GABA+ estimates for spant (up to 6 Hz) and Tarquin (2-6 Hz) were slightly moderated when separated from SNR effects; conversely, LCModel (over 5 Hz) and Gannet (Gaussian LB) showed greater changes when SNR effects were removed.

**Figure 6.**
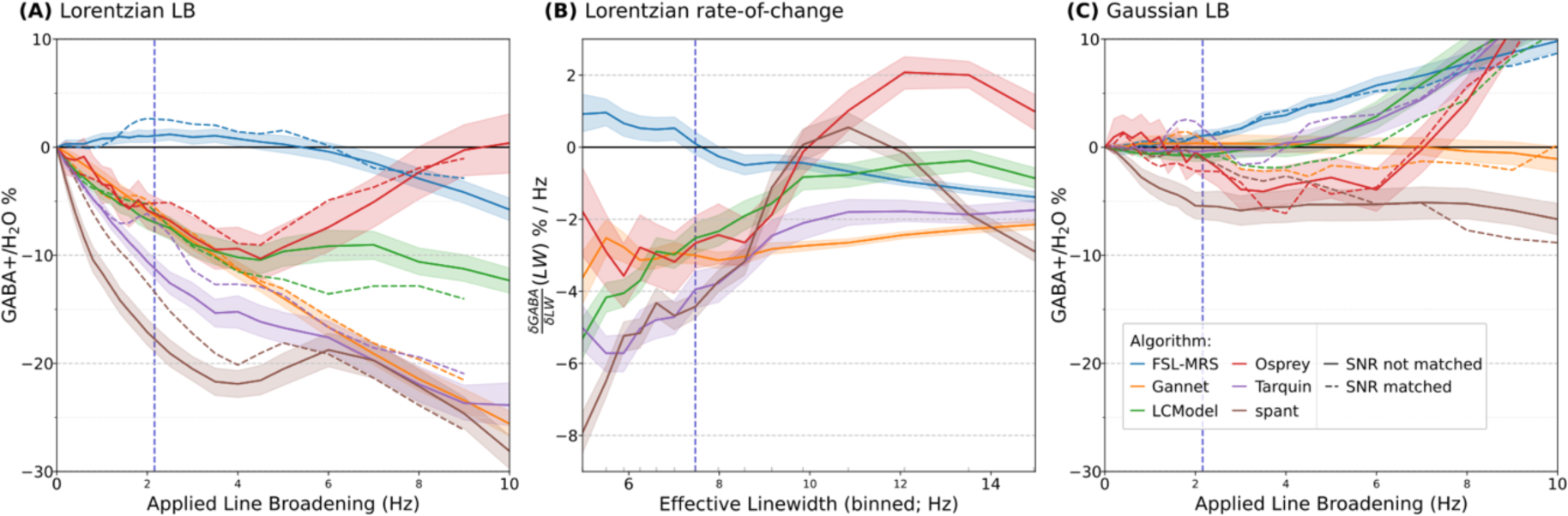
Showing the relative obtained GABA+ estimate as a function of applied Lorenzian (A) and Gaussian (C) linebroadening, and (B) the piecewise linear rate-of-change in GABA+ estimate for the Lorenzian case. Dashed vertical line indicates approximate range of the LLWF metric evaluation.

The LLWF metric (defined in section 2.5) is illustrated in Figure 6B, as the per-subject median 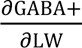 (LW) over the first 2.12 Hz, and indicates the degree to which estimates from a given algorithm are likely to be affected by a moderate change in linewidth. A comparison of this metric across different algorithms (grouped by vendor) is presented in Supplementary Figure 6, with numerical and statistical outcomes (unpaired t-tests) examining algorithm-specific effects in Supplementary Table 5, and vendor-specific effects (within algorithm) in Supplementary Table 6.

While most algorithms presented a similar general trend (lower GABA+ estimates as linewidth increased), the magnitude varied significantly (p_holm_ < 0.01) between all pairs of algorithms, except between LCModel, Gannet and Osprey and between spant and Tarquin, which happened to behave similarly in the defined range. Although the present analysis and discussion is focussed on GABA+, the interested reader may find similar plots for Glx_diff_, tNAA_diff_, and tCr_off_ in Supplementary Figure 7.

### 3.2 Synthetic Data

Quantification outcomes from the synthetic data are presented in Figure 7 for the GABA-only case and for the complete model (G and G+MBR respectively), with outcomes from intermediate model steps presented in Supplementary Figure 8, and fits in Supplementary Figure 9. Remarkably, the basic trend towards lower estimates at higher Lorentzian linewidth presents already in the trivial GABA-only case for most algorithms, although LCModel in particular maintained a flat response up to 7.5 Hz LW.

**Figure 7.**
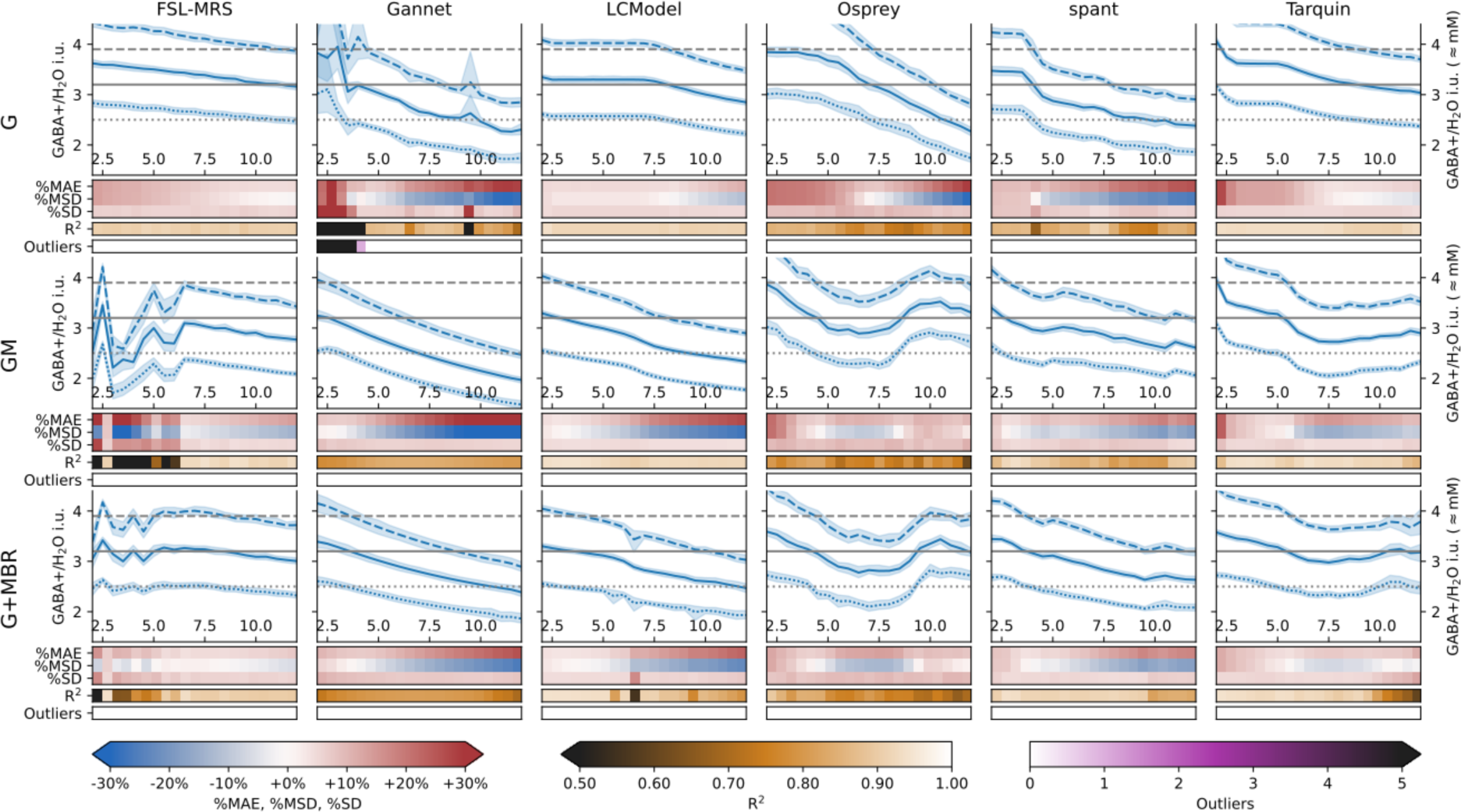
Quantification outcomes for each algorithm, modelling synthetic data of varying complexity: GABA only (G), GABA and additional metabolite components (GM), GABA+ with additional metabolite components and background signal (G+MBR).

FSL-MRS exhibited some variability in behaviour at lower linewidths in the intermediate model steps. This was largely (but not entirely) resolved in the full (G+MBR) model, and is explored further in Supplementary Section F (Supplementary Table 8 and Supplementary Figures 11, 12 and 13). Gannet also exhibited greater variability at lower (<5 Hz) linewidths, particularly for the lower simulated concentration – most likely an interaction between the GABA+ and baseline models. Once again, this variability was not observed with the full synthetic model. We note that Gannet by default applies 3 Hz linebroadening on top of the data’s inherent linewidth, so this is unlikely to affect in-vivo data in general usage.

Some algorithms, most notably Osprey, show distinct inflection points after stepping from the GABA-only synthetic data (G) to the GM model. The response of the Osprey model turns after 8 Hz, which reflects the behaviour seen in-vivo (see Figure 6). Tarquin exhibited a steep slope between 5-6 Hz, but was virtually flat between 7 and 11 Hz. These features became apparent only after stepping from the trivial GABA-only synthetic data (G) to data with other metabolites included (GM). LCModel presented a slight deviation at 6 Hz linewidth, most visible in the lower nominal GABA case; this is also reflected in a markedly reduced R^2^ at that point.

For most algorithms, error metrics %MAE and %MSD degrade progressively with linewidth, albeit to differing degrees. However, the %SD of estimates were not seen to degrade appreciably over this range, and most algorithms maintained their sensitivity to the nominal differences (assessed by R^2^) for a broad range of linewidths. Apparent variability in estimates constrained the sensitivity of FSL-MRS at lower linewidths.

## 4 Discussion and Conclusions

### 4.1 Limitations

#### 4.1.1 Scope and generalisability

While many of the key findings in this study present across several of the algorithms examined, they do not necessarily generalise further. A number of other algorithms and implementations are available to the MRS community^62–71^, each potentially differing in modelling strategy, baseline handling and constraint models and therefore likely to present unique characteristics. Most implementations are also highly configurable, with even subtle differences in modelling parameters or prior knowledge having potentially substantial impact on the behaviour, as observed in previous work^23,44^ and also here in the supplementary analyses (supplementary sections E, F, G). Since the configurations tested herein generally remain close to the developer-supplied default or recommended configurations, our findings are more representative of “typical use”, rather than the best achievable results with extensive local optimisation.

#### 4.1.2 In-vivo limitations

The in-vivo analysis is constrained by the absence of a ground truth; it is not possible to assess the absolute accuracy of measurements, only outcomes relative to the higher-quality unfiltered spectra. Taking random subsets of transients for the reduced SNR cases may over-estimate variance relative to contiguous acquisition of the same number of transients, since the longer scan may be subject to greater confounds associated with subject motion, thermal drift^72^, and perhaps real physiological variations.

The linebroadening operations applied to in-vivo data entail broadening (smoothing) across the entire frequency range, which also affects SNR. It is not straightforward to disentangle these factors. A common approach is to add noise back into the broadened spectra – however, while simple additive noise (Gaussian white noise for example) may yield similar SNR estimates, the characteristics of such noise are not necessarily representative of a true spectrum acquired with lower SNR. In the present analysis we instead separate these factors by numeric modelling, interpolating across a 2D grid of SNR and LB factors to estimate variation associated with each factor independently (see section 2.5). Nonetheless, the independence of noise samples contributing to the fixed-SNR estimates may be reduced on account of the linebroadening.

#### 4.1.3 Synthetic Data limitations

The lack of ground truth in the in-vivo analysis is addressed by additional analyses using carefully-prepared synthetic data^73^. However, any study involving simulated data is limited by the knowledge on which the simulation was based, and on simplifications and assumptions inherent in the simulation model – often the same knowledge which guides design of the modelling algorithms. The present study is no exception: the simulated metabolite signals come from the same basis set with which the data was subsequently fit. While we attempt to mitigate any gaps in our prior knowledge basis sets through incorporation of residual signals derived from in-vivo data fits, the simulation is nonetheless limited in other aspects, including potential asymmetric/non-Lorentzian/non-Gaussian lineshape changes, shifts in frequency or phase and other factors which may be expected to arise in in-vivo data.

### 4.2 Macromolecule Fitting, Baseline and Artefact Rejection around 3.2 ppm

Previous studies have identified a number of factors potentially affecting the accuracy of quantification of GABA-edited MRS data and worthy of further attention. Key amongst these is the handling of co-edited macromolecule and metabolite signal underlying the 3.0 ppm GABA peak ^44,74–78^. Although broadly targeted in the simulation study, the reliability of estimates from simulated data was not substantially degraded by inclusion of this component in the simulation model, suggesting that the fitting models as configured perform reasonably well at modelling the aggregate signal (GABA+). Similarly, other baseline distortions in the data were not found to drive the linewidth-related bias. This is compatible with recent findings on simulated PRESS data, where most of the packages considered performed equally well on datasets with and without macromolecules^17^. We emphasize that the current study looks only at GABA+, and studies attempting to quantify GABA alone are likely to be affected differently.

Another commonly observed artefactual feature in GABA-edited MEGA-PRESS spectra is a small hump around 3.2 ppm, which may arise due to incomplete subtraction of choline ^79^, or contributions from other co-edited signals (perhaps including valine-containing macromolecules ^80^ and arginine ^81^). Although not explicitly modelled in this study, this component is subtly present in the consensus background signals applied in the simulation study. Once again, it does not appear to drive the linewidth-related effects observed here.

Although these factors were not found to be significant specifically in relation to the linewidth-related bias, they remain items of interest in further refinement of fitting algorithms.

### 4.3 Factors contributing to the observed changes

The simulation analysis described in section 2.3 was designed to identify specific features giving rise to the observed effects in the in-vivo data. Surprisingly, several algorithms presented a linewidth-related effect even for the most trivial case of a single metabolite signal on a flat background. This suggests that some of the observed effects relate to the basic lineshape model, or are intrinsic to the optimisation techniques applied and not solely dependent on any particular confounding signal. This may indicate that modelling optimization is not sufficiently responsive to changing linewidth, instead preferring to ascribe some part of the signal to other model components (perhaps including the baseline model). It is possible that further tuning of cost factors associated with linebroadening may mitigate this effect.

The influence of lineshape model should not be underestimated; while basis set peaks and applied linebroadening herein exhibit a basic Lorentzian lineshape, measured in-vivo metabolite peaks will present a more complex profile – closer to a Voigt profile^45^. Discrepancies between the real and modelled lineshape have been shown to introduce systematic errors in quantification^3,4^. This may be a contributing factor particularly for algorithms having only a basic lineshape model (e.g., Gannet which by default uses a Gaussian fit). Even where a more nuanced lineshape model is used, this is often dominated by a small number of major peaks (such as Cho and Cr for short TE PRESS, or NAA for GABA-edited data. It is not certain that an identical model will be applicable to all peaks – particularly when transferring from strong singlets to complex, edited multiplets. Hence, a mismatch in lineshape between the major peaks dominating the lineshape model, and the peaks to which that model is applied, may account for some of the variation seen herein. This may be mitigated by allowing refined per-peak lineshape modelling – although such modelling may be less robust to noise in the case of smaller peaks. Indeed, the exploratory analysis for FSL-MRS (supplementary section F) showed, for that algorithm, having the same lineshape and shift parameters across all metabolite components (including the strong negative NAA peak) appeared to mitigate linewidth-related changes in GABA+ estimate. In practice, a well-tuned, firm but not rigid constraint may prove most suitable.

Another major factor differentiating the algorithms is baseline parameterization. LCModel, Osprey and spant all adopt a spline baseline (with adaptive spacing in the case of spant); in each of these cases, the baseline in higher linewidth cases is seen to be pulled substantially upwards (see Figure 3), through the apparent GABA peak – likely contributing to the underestimation in those cases. Simpler baseline models, such as the polynomial baseline of FSL-MRS, or Gannet’s sinusoidal baseline model, appear more robust in this regard – although the Gannet implementation becomes problematic in low SNR cases, leading to higher rejection rates.

### 4.4 Accounting for lineshape

For several of the algorithms tested, even a small difference in Lorentzian linewidth (0.5 Hz) between otherwise identical datasets could yield a statistically significant difference in obtained concentrations; this is concerning when considering comparisons across different regions, or between subject groups where spectral quality may be expected to differ, even slightly. Somewhat counterintuitively, the risk of false positive outcomes actually increases with “better” datasets: higher SNR data is likely to give reduced overall variance, and larger subject groups increase the statistical power to detect these subtle differences. With different metabolites exhibiting differing degrees of variation according to linewidth (see Supplementary Figure 7), scaling to an internal concentration reference (such as tCr_off_) subject to similar linewidth changes is not likely to be effective at mitigating these effects.

Since Lorentzian and Gaussian linebroadening factors exhibit contrasting behaviours, basic linewidth matching with a pure Lorentzian function (as frequently seen in the fMRS literature) may not be effective if this does not reflect the shape of the effects to be corrected, and may in fact exacerbate quantitative differences. If linewidth matching is to be performed, it is essential that an appropriate broadening function be chosen. If the nature of prospective differences is unknown, line*shape* matching (for example, by a deconvolution approach) may be more appropriate. We note however that matching approaches have limitations and complexities, some of which may be addressed by matching individual basis set components^82^ rather than the complete metabolite spectrum.

A more generalisable approach would be to routinely consider estimated linewidth as a covariate within statistical analysis of metabolite estimates, in any situation where signal quality may vary between groups. However, this may give rise to an unacceptable loss of statistical power, especially in cases where the linewidth correlates strongly with other (real) measurable effects.

### 4.5 Implications for Quality Control

Although the overall accuracy of estimates (in terms of %MAE, %MSD) deteriorated with increasing linewidth for most algorithms, the sensitivity to changes in concentrations (as reflected by %SD and moreover R^2^) was largely maintained across the range of linewidths assessed in the synthetic datasets; therefore, within the investigated range, accounting for lineshape as described in section 4.4 may be more advantageous than strict rejection criteria based on linewidth.

Similarly, the in-vivo study showed no strong group-level effects associated with SNR, except in high Gaussian LB cases, suggesting that estimates from lower-SNR data may remain usable in certain contexts^18^. We therefore re-iterate the need for quality control procedures which look beyond these basic signal metrics: close attention to the shape of the data, fit and residuals, and to the distribution of estimates themselves, may be more meaningful (and perhaps less prone to bias) than strict thresholding based on these metrics alone.

### 4.6 Key Recommendations

Based on our findings, we make the following recommendations:

- In any contexts where acquired spectral linewidth may vary between experimental groups, this must be rigorously controlled.
- Linewidth matching may be problematic if the broadening function applied does not accurately reflect the broadening mechanisms in the data; a more refined line*shape* matching approach may be preferable.
- Inclusion of linewidth as a covariate in statistical modelling may be more generalisable, and avoids pitfalls associated with matching.
- Controlling for differences in linewidth should be prioritised over simply rejecting datasets with linewidth over a certain threshold.

Additionally, we propose two areas of interest for potential refinement of these modelling algorithms:

- Broadly, further development of modelling algorithms should pay close attention to cost factors associated with linewidth, and how the model optimization may behave with respect to changes in linewidth of the source data.
- More specifically, revisiting the manner in which the lineshape model is derived, and whether this is appropriate to all the peaks being modelled. Lineshape variation is likely to differ between metabolite and macromolecule components.

While the effects described herein present across multiple tested configurations, the exact extent can vary significantly – we therefore re-iterate the need for consistency in modelling and reporting, and the value of converging on consensus best practices in this regard^21^.

### 4.7 Conclusions

We show that several contemporary fitting algorithms exhibit strong associations between linewidth and concentration estimate in response to LB_lorentz_ (T2 differences), and more moderate, contrasting associations in relation to LB_gauss_ (T2*/inhomogeneity effects).

The finding of reduced concentration estimates as a function of linewidth is consistent with much earlier findings^15,16^ for short TE STEAM and simulated datasets, where similar trends may be observed for several metabolites in certain conditions, using Hankel-Lanczos singular value decomposition (HLSVD)^83,84^, LCModel and AMARES algorithms for modelling. This is also consistent with more recent findings^17^ for major metabolites in simulated PRESS data, for several algorithms. That these trends persist after more than twenty years of algorithmic evolution leads to two conclusions: first, such patterns deserve further investigation and concerted effort to mitigate within the design of the modelling algorithms. At the same time, such trends must not be overlooked by users of the algorithms in analysing and interpreting their results.

Our findings highlight the ongoing need for rigorous consideration of potential linewidth differences between samples or conditions – and furthermore highlight the importance of adopting appropriate strategies to account for this.

## Acknowledgements

Analysis was performed within a project funded by ERC grant #249516, which additionally supported the contributions of ARC, LE, KH. Data used in this analysis were previously collected through an international collaborative study funded under NIH grant R01 EB016089. TKB acknowledges support from a One Child Every Child Postdoctoral Fellowship and a Cumming School of Medicine Postdoctoral Fellowship. ADH holds a Canada Research Chair in Magnetic Resonance Spectroscopy in Brain Injury and acknowledges funding support from the Natural Sciences and Engineering Research Council (NSERC) of Canada. GO acknowledges funding support from R00 AG062230, R21 EB033516, and P41 EB031771.

## 5 Declaration of Interest

The authors declare no conflicting interests.

## 6 Data Availability Statement

Scripts used for the present analysis are publicly available here; further dependencies are described within: https://git.app.uib.no/bergen-fmri/gaba-linewidth

Spectra analysed in this manuscript were obtained from the publicly available Big GABA repository on NITRC, https://www.nitrc.org/projects/biggaba

Basis sets used in the primary analysis were obtained from the publicly available Osprey package, https://schorschinho.github.io/osprey

## List of Abbreviations

BOLD: *blood oxygen level dependent*
Cho: *Choline*
Cr: *Creatine* diff *difference (edited) spectrum*
FD: *frequency domain*
FID: *Free Induction Decay (time-domain MRS signal)*
FWHM: *linewidth full width at half maximum*
G(+)[MBR]: *Synthetic spectral models, see section 2.3*
GABA: *γ-aminobutyric acid*
GABA+: *total edited signal at 3 ppm; GABA with underlying coedited signals*
Gln: *Glutamine*
Glu: *Glutamate*
GPC: *glycerophosphorylcholine*
GSH: *glutathione*
HLSVD: *Hankel-Lanczos singular value decomposition*
i.u.: *institutional units*
Lac: *Lactate*
LBgauss: *Gaussian linebroadening factor*
LBlorentz: *Lorentzian linebroadening factor*
LCM: *linear combination modelling*
LLWF: *Linear line-width factor, see section 2.5*
MAD: *Median Absolute Deviation*
MAE: *Mean Absolute Error*
MEGA-PRESS: *Mescher–Garwood point-resolved spectroscopy*
mI: *myo-inositol*
MRS: *Magnetic Resonance Spectroscopy*
MSD: *Mean Signed Difference*
NAA: *N-acetyl aspartate*
NAAdiff: *NAA measured from the difference spectrum*
NAAG: n-acetylaspartylglutamate
PCh: *phosphorylcholine*
PCr: *phosphocreatine*
pholm: *Holm-Bonferroni adjusted p-value*
punc: *Uncorrected p-value*
R1-3: *adopted rejection criteria; see methods section 2.1.2*
SD: *Standard Deviation*
SNR: *Signal-to-Noise Ratio*
T2*: *Effective transverse relaxation rate*
tCroff: *total creatine measured from the edit-OFF sub-spectrum*
TD: *time domain*
VPC: *Variance Partition Coefficients*

## Notes

### Competing Interest Statement

The authors have declared no competing interest.

