## Supplementary Material for "Linewidth-related bias in modelled concentration estimates from GABA-edited ^1^H-MRS"

A MRSinMRS checklist ... 2
B Acquisition Details ... 3
C Supplementary Details: In-vivo data ... 4
D Supplementary Details: Synthetic Data ... 10
D.1 Synthetic data generation ... 10
D.2 Outcomes from fits to synthetic data ... 11
E Exploratory Analysis #1: spant baseline modelling ... 13
E.1 Background and Methods ... 13
E.2 Results and Discussion ... 14
F Exploratory Analysis #2: FSL-MRS baseline modelling, optimization and lineshape ... 17
F.1 Background and Methods ... 17
F.2 Results and Discussion ... 18
G Exploratory Analysis #3: LCModel baseline knot spacing ... 22
G.1 Background and Methods ... 22
G.2 Results and Discussion ... 22
H Supplementary Methods ... 24
H.1 Difference spectrum modelling ... 24
H.1.1 FSL-MRS ... 24
H.1.2 Gannet ... 25
H.1.3 LCModel ... 25
H.1.4 Osprey ... 26
H.1.5 spant ... 26
H.1.6 Tarquin ... 27
H.2 Creatine Reference: Edit-OFF sub-spectrum modelling ... 28
H.2.1 FSL-MRS ... 28
H.2.2 Gannet ... 29
H.2.3 LCModel ... 29
H.2.4 Osprey ... 29
H.2.5 spant ... 29
H.2.6 Tarquin ... 30

### A MRSinMRS checklist

| | |
|---|---|
| Site (name or number) | [GPS](1-10) |
| **1. Hardware** | |
| a. Field strength [T] | 3 T |
| b. Manufacturer | See sup. table 2 |
| c. Model (software version if available) | |
| d. RF coils: nuclei (transmit/receive), number of channels, type, body part | |
| e. Additional hardware | N/A |
| **2. Acquisition** | |
| a. Pulse sequence | MEGA-PRESS (GABA+ editing; see sup. table 2 for variants) |
| b. Volume of interest (VOI) locations | Medial parietal lobe |
| c. Nominal VOI size [cm3,mm3] | 30 x 30 x 30 mm3 |
| d. Repetition time ($T_R$), echo time ($T_E$) [ms, s] | $T_R$ = 2000 ms, $T_E$ = 68 ms |
| e. Total number of excitations or acquisitions per spectrum<br>In time series for kinetic studies<br>i. Number of averaged spectra (NA) per time point<br>ii. Averaging method (eg block-wise or moving average)<br>iii. Total number of spectra (acquired/in time series) | Up to 320 averages (160 ON and 160 OFF transients) |
| f. Additional sequence parameters (spectral width in Hz, number of spectral points, frequency offsets)<br>If STEAM:, mixing time ($T_M$)<br>If MRSI: 2D or 3D, FOV in all directions, matrix size, acceleration factors, sampling method | See sup. table 2 for spectral width and number of points.<br>15 ms editing pulses at 1.9 ppm (ON) and 7.46 ppm (OFF) |
| g. Water suppression method | See sub. table 2 |
| h. Shimming method, reference peak, and thresholds for “acceptance of shim” chosen | See sup. table 2; acceptance criteria varied across sites |
| i. Triggering or motion correction method (respiratory, peripheral, cardiac triggering, incl. device used and delays) | N/A |
| **3. Data analysis methods and outputs** | |
| a. Analysis software | FSL-MRS 1.1.1<br>Gannet 3.1<br>Osprey 1.0.1.1<br>LCModel 6.3-1P<br>spant 1.13.9<br>Tarquin 4.3.11 (local patch) |
| b. Processing steps deviating from quoted reference or product | Filtering to experimentally adjust SNR and linewidth, see section 2.1.1 |
| c. Output measure (eg absolute concentration, institutional units, ratio), processing steps deviating from quoted reference or product | Water-referenced estimates (with adjustment for tissue content); see section 2.4 |
| d. Quantification references and assumptions, fitting model assumptions | Standardised basis set where applicable, incorporating 3.0 ppm macromolecule component; see sections 2.2, 2.4 |
| **4. Data quality** | |
| a. Reported variables (SNR, linewidth (with reference peaks)) | SNR and Linewidth from $NAA_{diff}$ |
| b. Data exclusion criteria | R1: Correlation with normalized mean spectrum < 0.5 across metabolite range, or deviating from group mean by > 3 SD<br>R2: SNR < 80 ($NAA_{diff}$) or FWHM linewidth > 10 Hz ($NAA_{diff}$)<br>See section 2.1.2 |
| c. Quality measures of postprocessing model fitting (eg CRLB, goodness of fit, SD of residual) | R3: Strong outliers (> 5 x median absolute deviation), see 2.1.2<br>%SD across variants based on same dataset;<br>% mean signed and abs. errors (%MSE, %MAE) for synthetic data |
| d. Sample spectrum | See Sup Figure 1 |

*Supplementary Table 1: MRSinMRS checklist[1] summarising key details of the MRS acquisition*

### B Acquisition Details

| Site | N | Age (years±SD) | Sex F / M | Scanner vendor and model | Software release | Tx/Rx hardware | MEGA-PRESS sequence variant | Phase cycling | Editing inter-leaving | B0 shimming approach | Water suppr. | Spectral width (Hz) | Data points |
|---|---|---|---|---|---|---|---|---|---|---|---|---|---|
| MRSinMRS | | | | 1b,c | 1c | 1d | 2a | | | 2h | 2g | 2f | 2f |
| G1 | 7 | 22.9±3.7 | 4 / 3 | GE Discovery MR750w | DV25 | Body coil/32-ch head coil | Interleaved sequence | 2 | 2 | Double-echo GRE | CHESS | 5000 | 4096 |
| G4 | 12 | 25.6±4.5 | 6 / 6 | GE Discovery MR750 | DV25 | Body coil/8-ch head coil | ATSM patch | 8 | 1 | Double-echo GRE | CHESS | 5000 | 4096 |
| G5 | 12 | 25.5±3.7 | 5 / 7 | GE Discovery MR750 | DV25 | Body coil/32-ch head coil | ATSM patch | 8 | 1 | Double-echo GRE | CHESS | 2000 | 2048 |
| G6 | 12 | 24.3±4.2 | 6 / 6 | GE Signa HDx | HD16 | Body coil/8-ch head coil | ATSM patch | 2 | 2 | Double-echo GRE | CHESS | 2000 | 2048 |
| G7 | 12 | 28.1±4.0 | 6 / 6 | GE Discovery MR750 | DV24 | Body coil/8-ch head coil | ATSM patch | 8 | 1 | Double-echo GRE | CHESS | 2000 | 2048 |
| G8 | 12 | 29.7±2.1 | 6 / 6 | GE Discovery MR750 | DV24 | Body coil/8-ch head coil | ATSM patch | 8 | 1 | Double-echo GRE | CHESS | 2000 | 2048 |
| **All G** | **67** | **26.2±4.3** | **33 / 34** | | | | | | | | | | |
| P1 | 9 | 25.0±3.7 | 4 / 5 | Philips Achieva | R5.1.7 | Body coil/32-ch head coil | JHU patch | 16 | 1 | PB-auto | VAPOR | 2000 | 2048 |
| P3 | 12 | 25.1±2.9 | 6 / 6 | Philips Achieva | R3.2.2 | Body coil/32-ch head coil | JHU patch | 16 | 1 | PB-auto | VAPOR | 2000 | 2048 |
| P4 | 12 | 29.2±3.1 | 5 / 7 | Philips Ingenia CX | R5.1.7 | Body coil/32-ch head coil | JHU patch | 16 | 1 | PB-auto | MOIST | 2000 | 2048 |
| P5 | 12 | 24.9±4.3 | 7 / 5 | Philips Achieva TX | R5.1.7 | Body coil/32-ch head coil | JHU patch | 16 | 1 | PB-auto | MOIST | 2000 | 2048 |
| P6 | 8 | 23.1±2.4 | 3 / 5 | Philips Achieva | R3.2.3 | Body coil/8-ch head coil | JHU patch | 16 | 1 | PB-auto | MOIST | 2000 | 2048 |
| P7 | 12 | 27.3±3.7 | 7 / 5 | Philips Ingenia | R5.1.8 | Body coil/32-ch head coil | JHU patch | 16 | 1 | PB-auto | VAPOR | 2000 | 2048 |
| P8 | 12 | 23.6±3.7 | 6 / 6 | Philips Ingenia CX | R5.1.8 | Body coil/32-ch head coil | JHU patch | 16 | 1 | PB-auto | MOIST | 2000 | 2048 |
| P9 | 12 | 23.2±2.0 | 5 / 7 | Philips Achieva | R5.1.7 | Body coil/32-ch head coil | JHU patch | 16 | 1 | PB-auto | VAPOR | 2000 | 2048 |
| P10 | 12 | 25.8±4.6 | 6 / 6 | Philips Ingenia | R5.1.9 | Body coil/15-ch head coil | JHU patch | 16 | | PB-auto | MOIST | 2000 | 2048 |
| **All P** | **101** | **25.4±3.9** | **49 / 52** | | | | | | | | | | |
| S1 | 12 | 25.7±3.7 | 6 / 6 | Siemens Trio | VB17 | Body coil/32-ch head coil | WIP (529) | 16 | 1 | 3D-DESS + manual | CHESS | 4000 | 4096 |
| S3 | 12 | 31.6±3.4 | 9 / 3 | Siemens Prisma | VD13 | Body coil/20-ch head/neck coil | WIP (859D) | 16 | 1 | FAST(EST) MAP | WET | 4000 | 4096 |
| S5 | 12 | 26.5±3.7 | 6 / 6 | Siemens Trio | VB17 | Body coil/12-ch head coil | WIP (529) | 16 | 1 | 3D-DESS | CHESS | 4000 | 4096 |
| S6 | 6 | 26.2±2.0 | 1 / 5 | Siemens Trio | VB17 | Body coil/32-ch head coil | WIP (529) | 16 | 1 | FAST(EST) MAP | WET | 4000 | 4096 |
| S8 | 12 | 24.0±3.5 | 11 / 1 | Siemens Prisma | VE11 | Body coil/64-ch head coil | WIP (859G) | 16 | | 3D-DESS | WET | 4000 | 4096 |
| **All S** | **54** | **26.9±4.3** | **33 / 21** | | | | | | | | | | |
| **Total** | **222** | **26.0±4.1** | **115 / 107** | | | | | | | | | | |

*Supplementary Table 2: Basic demographics, hardware and software parameters for the constituent datasets; reproduced from previously published work [2].*

### C Supplementary Details: In-vivo data

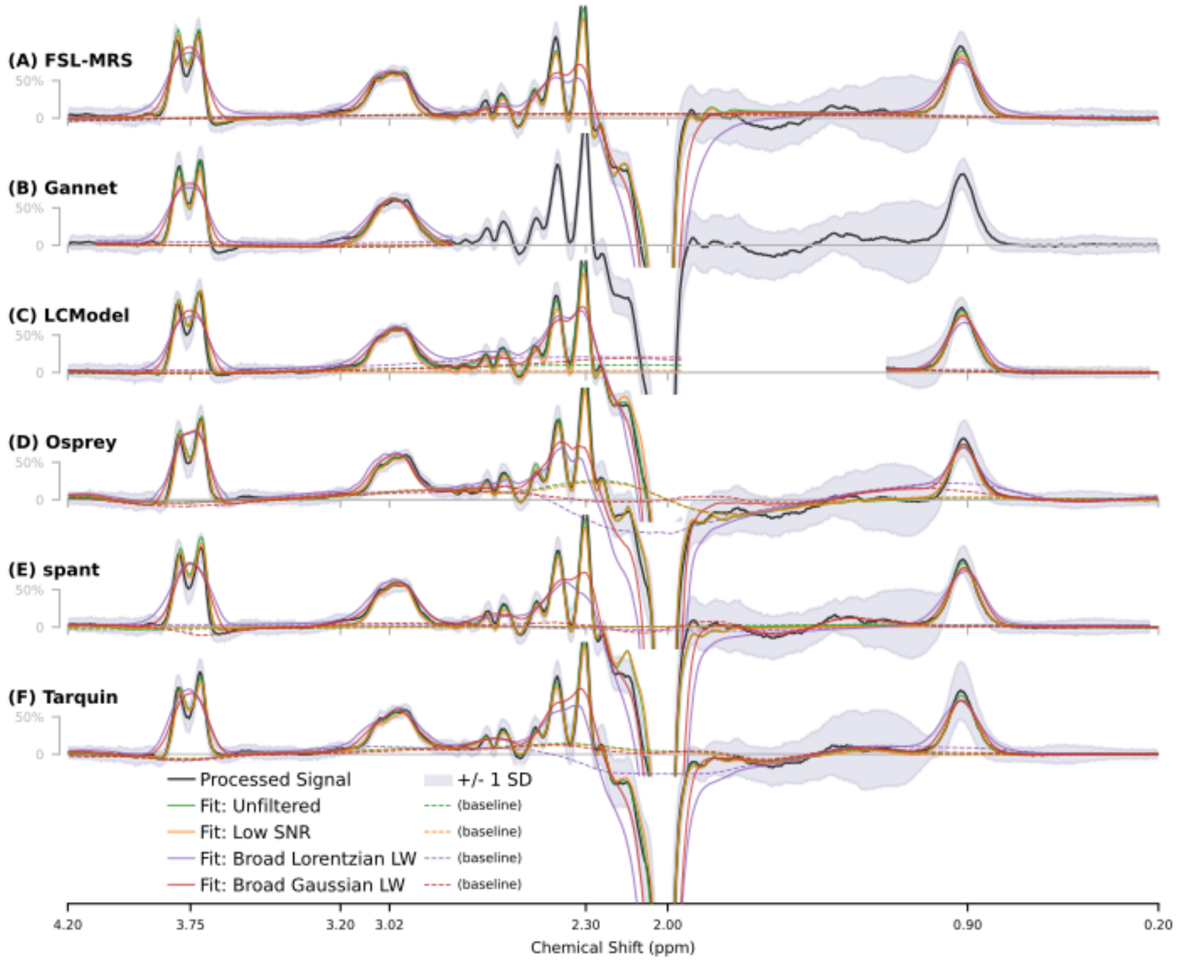


*Supplementary Figure 1 Group-average fit by each algorithm, to original (unfiltered) data, low SNR and high linewidth variants*

| Site | SNR | Linewidth (Hz) | Site | SNR | Linewidth (Hz) |
|---|---|---|---|---|---|
| G1 | 142 [124; 206] | 5.64 [5.11; 7.33] | P1 | 222 [150; 306] | 4.71 [4.01; 5.59] |
| G4 | 133 [104; 156] | 6.74 [5.22; 7.86] | P10 | 104 [85.9; 129] | 5.65 [4.32; 6.92] |
| G5 | 167 [118; 193] | 5.91 [4.64; 7.27] | P3 | 142 [109; 176] | 5.00 [4.21; 6.3] |
| G6 | 141 [127; 188] | 5.46 [4.72; 6.53] | P4 | 242 [202; 273] | 4.50 [4.11; 4.9] |
| G7 | 133 [97.2; 159] | 6.19 [5.51; 8.19] | P5 | 196 [168; 271] | 5.00 [4.27; 5.55] |
| G8 | 120 [82.9; 150] | 5.59 [4.73; 6.61] | P6 | 151 [102; 193] | 4.62 [4.39; 6.01] |
| S1 | 221 [165; 305] | 6.17 [4.32; 8.77] | P7 | 175 [155; 199] | 7.61 [5.81; 8.87] |
| S3 | 160 [84.3; 232] | 4.97 [4.17; 5.81] | P8 | 257 [189; 308] | 4.71 [3.69; 5.97] |
| S5 | 166 [121; 193] | 5.52 [4.76; 9.41] | P9 | 216 [172; 255] | 4.69 [4.05; 5.18] |
| S6 | 271 [190; 289] | 5.59 [4.41; 5.95] | | | |
| S8 | 240 [167; 264] | 4.49 [4.2; 4.78] | **All** | **171 [104; 273]** | **5.33 [4.25; 7.61]** |

*Supplementary Table 3 : Basic quality metrics for incoming data (before filtering), per site – expressed as median [5, 95]-percentiles, derived from tNAA measured on the difference spectrum.*

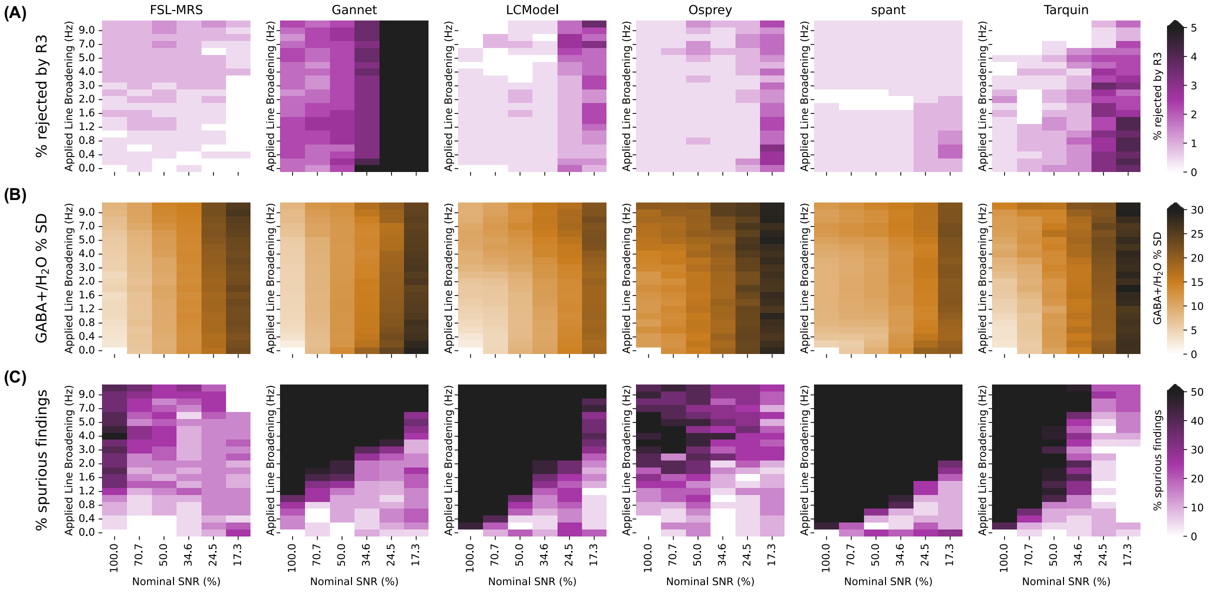


*Supplementary Figure 2 : Rejection rate according to R3 (A), standard deviation of GABA+ estimates (B), and rate of "spurious findings": false-positive statistically significant difference in mean outcome, on a per-site basis (C), given the applied change in SNR/LW for **Lorentzian** linebroadening*

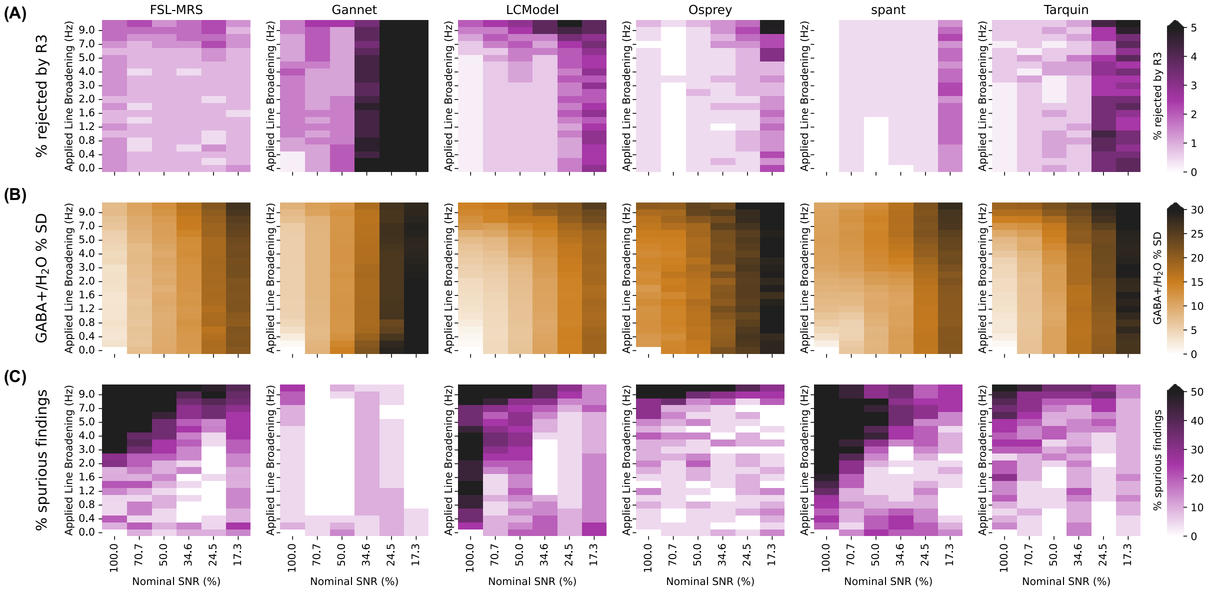


*Supplementary Figure 3 Rejection rate according to R3 (A), standard deviation of GABA+ estimates (B), and rate of "spurious findings": false-positive statistically significant difference in mean outcome, on a per-site basis (C), given the applied change in SNR/LW for **Gaussian** linebroadening*

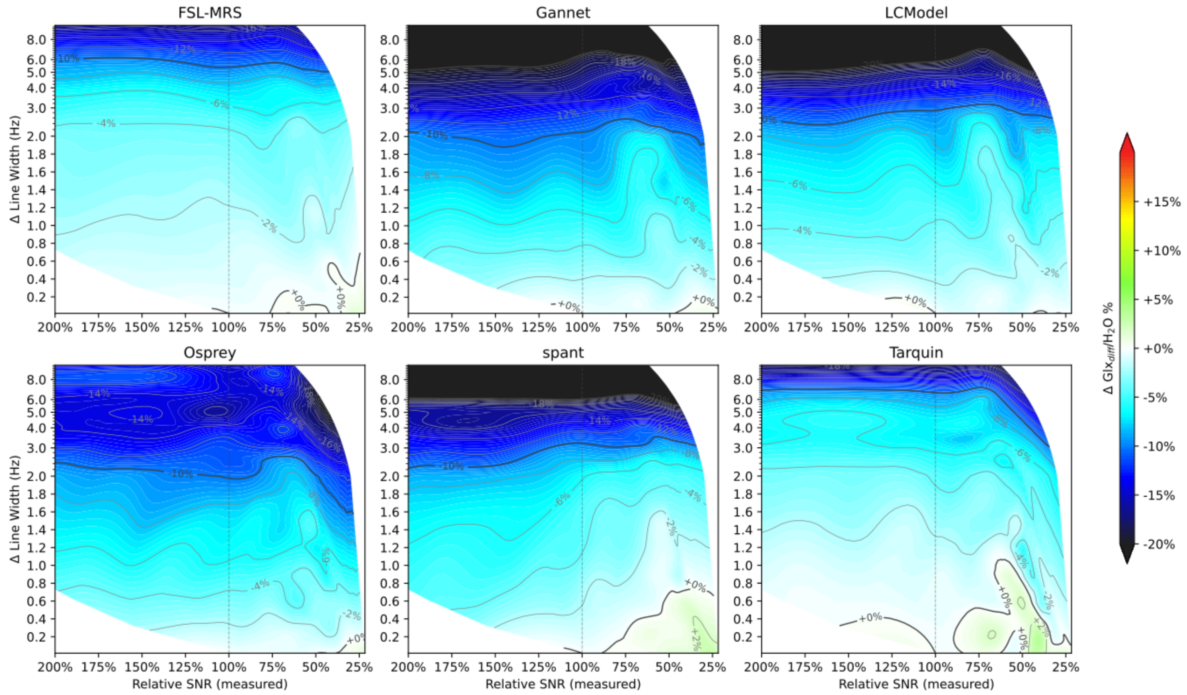


*Supplementary Figure 4 Relative Glx estimate as a function of Lorentzian linebroadening and SNR, for each of the algorithms assessed*

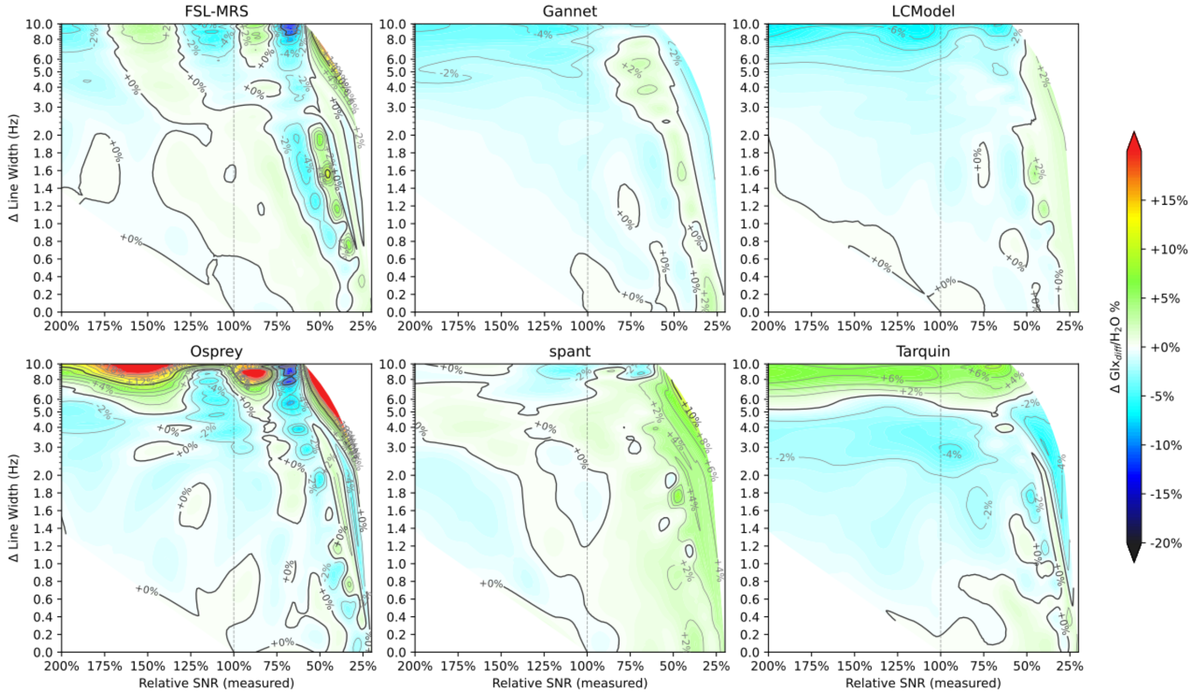


*Supplementary Figure 5 Relative Glx estimate as a function of Gaussian linebroadening and SNR, for each of the algorithms assessed*

| | | Lorentzian LB | | | | | | | | | | | Gaussian LB | | | | | | | | | | |
|---|---|---|---|---|---|---|---|---|---|---|---|---|---|---|---|---|---|---|---|---|---|---|---|
| Line Broadening | | | | | | | | 0 Hz | 3 Hz | 10 Hz | all | | | | | | | | 0 Hz | 3 Hz | 10 Hz | all | |
| Relative SNR | | 100% | 71% | 50% | 35% | 24% | 17% | | | | all | | 100% | 71% | 50% | 35% | 24% | 17% | | | | all | |
| Algorithm | VPC | | | | | | | | | | | Median | | | | | | | | | | | Median |
| FSL-MRS | Subject | 87,3 | 89 | 91,3 | 92,5 | 93,5 | 90,9 | 31,5 | 33,6 | 38,3 | 40,4 | 88,15 | 84,5 | 88,3 | 88,1 | 90,5 | 92,3 | 93,8 | 34,9 | 35,1 | 38,4 | 45,2 | 86,3 |
| | LB | 2,2 | 1,6 | 1,4 | 1 | 0,7 | 1,3 | | | | 1,1 | 1,3 | 6,9 | 5,5 | 5 | 3 | 1,7 | 1,2 | | | | 2,6 | 3 |
| | SNR | | | | | | | 3,2 | 0,2 | 0 | 0,5 | 0,35 | | | | | | | 2,2 | 0,3 | 0,1 | 0,6 | 0,45 |
| | Resid | 10,5 | 9,4 | 7,3 | 6,4 | 5,9 | 7,9 | 65,3 | 66,2 | 61,7 | 58 | 9,95 | 8,6 | 6,2 | 7 | 6,5 | 6 | 5 | 62,9 | 64,5 | 61,4 | 51,5 | 7,8 |
| Gannet | Subject | 55 | 61,8 | 73,5 | 74,5 | 77,9 | 84,4 | 20,1 | 24,7 | 28,5 | 28,6 | 58,4 | 92,9 | 94,1 | 92,8 | 93,1 | 91,3 | 91 | 16,2 | 21,8 | 26,9 | 32 | 91,15 |
| | LB | 39,4 | 30,6 | 21,2 | 17,7 | 12,4 | 6,6 | | | | 16,1 | 17,7 | 0,1 | 0 | 0 | 0 | 0,1 | 0,1 | | | | 0 | 0 |
| | SNR | | | | | | | 0 | 0 | 0 | 0,1 | 0 | | | | | | | 0 | 0 | 0 | 0,2 | 0 |
| | Resid | 5,6 | 7,6 | 5,3 | 7,9 | 9,6 | 9 | 79,9 | 75,3 | 71,5 | 55,2 | 9,3 | 7 | 5,9 | 7,2 | 6,9 | 8,6 | 9 | 83,8 | 78,2 | 73,1 | 67,8 | 8,8 |
| LCModel | Subject | 60 | 63,4 | 69,2 | 69,7 | 70,1 | 77,2 | 28,6 | 36,5 | 37,2 | 30,2 | 61,7 | 58,1 | 63,3 | 66,3 | 74,3 | 84,7 | 83,7 | 35,2 | 32,2 | 42,8 | 34,3 | 60,7 |
| | LB | 10,7 | 10,5 | 12 | 12,6 | 12,9 | 9,8 | | | | 10,6 | 10,7 | 13 | 9,7 | 7,1 | 4,7 | 1,9 | 0,7 | | | | 3,9 | 4,7 |
| | SNR | | | | | | | 2,8 | 0,1 | 1,9 | 0,4 | 1,15 | | | | | | | 2 | 0,6 | 0 | 0,5 | 0,55 |
| | Resid | 29,3 | 26,1 | 18,8 | 17,7 | 17 | 13 | 68,6 | 63,3 | 60,9 | 58,8 | 27,7 | 28,9 | 27 | 26,6 | 21,1 | 13,4 | 15,6 | 62,8 | 67,2 | 57,2 | 61,2 | 27,95 |
| Osprey | Subject | 45,4 | 54,5 | 60,9 | 67,8 | 73,9 | 79,4 | 22,9 | 24,9 | 48,9 | 31,7 | 51,7 | 43,8 | 52,9 | 59,6 | 72 | 75 | 83,2 | 26,5 | 24,6 | 35,4 | 30,6 | 48,35 |
| | LB | 4,9 | 4,1 | 3,5 | 2 | 1,6 | 0,9 | | | | 2,2 | 2,2 | 9,9 | 8 | 6,3 | 3,8 | 3,1 | 1,1 | | | | 3,9 | 3,9 |
| | SNR | | | | | | | 0,1 | 0 | 0 | 0,3 | 0,05 | | | | | | | 0 | 0 | 0,7 | 0 | 0 |
| | Resid | 49,7 | 41,4 | 35,6 | 30,2 | 24,5 | 19,7 | 77 | 75,1 | 51,1 | 65,8 | 45,55 | 46,3 | 39,2 | 34,1 | 24,2 | 21,9 | 15,7 | 73,5 | 75,4 | 63,9 | 65,5 | 42,75 |
| spant | Subject | 43,2 | 41,2 | 42,9 | 45,9 | 48,6 | 57,4 | 34,9 | 30,9 | 56,6 | 21,6 | 43,05 | 70,3 | 64,8 | 67,7 | 66,2 | 67,6 | 73,4 | 38,2 | 35,6 | 33,8 | 39,2 | 65,5 |
| | LB | 31,5 | 34 | 32,8 | 30,3 | 25,4 | 19,8 | | | | 25,7 | 30,3 | 2,7 | 4,1 | 4,5 | 3,8 | 2,4 | 1,9 | | | | 2,5 | 2,7 |
| | SNR | | | | | | | 1,9 | 2,9 | 0 | 1,6 | 1,75 | | | | | | | 2,3 | 2,7 | 0,2 | 1,9 | 2,1 |
| | Resid | 25,3 | 24,8 | 24,3 | 23,8 | 26 | 22,9 | 63,1 | 66,2 | 43,4 | 51,1 | 25,65 | 27 | 31,1 | 27,8 | 30 | 30,1 | 24,7 | 59,5 | 61,8 | 66 | 56,4 | 30,6 |
| Tarquin | Subject | 56,4 | 62,2 | 70,5 | 70,7 | 76,2 | 81,2 | 22,5 | 29,6 | 41,8 | 30,9 | 59,3 | 68 | 72,4 | 79,9 | 83,4 | 84,7 | 88 | 32,4 | 32,4 | 54,2 | 43,6 | 70,2 |
| | LB | 24,9 | 21,2 | 15 | 14,5 | 10,7 | 6,9 | | | | 12,5 | 14,5 | 8,6 | 7 | 4,5 | 2,4 | 1,9 | 0,4 | | | | 2,4 | 2,4 |
| | SNR | | | | | | | 2,1 | 4,7 | 2 | 3,1 | 2,6 | | | | | | | 1,2 | 2,3 | 0 | 1,6 | 1,4 |
| | Resid | 18,7 | 16,7 | 14,5 | 14,8 | 13 | 12 | 75,4 | 65,7 | 56,2 | 53,5 | 17,7 | 23,4 | 20,6 | 15,6 | 14,2 | 13,3 | 11,6 | 66,3 | 65,4 | 45,8 | 52,3 | 22 |

*Supplementary Table 4 Variance Partition Coefficients (VPCs) for GABA+ estimated from filtered spectra at specified SNR/Line Broadening levels; all VPCs quoted as percentages.*

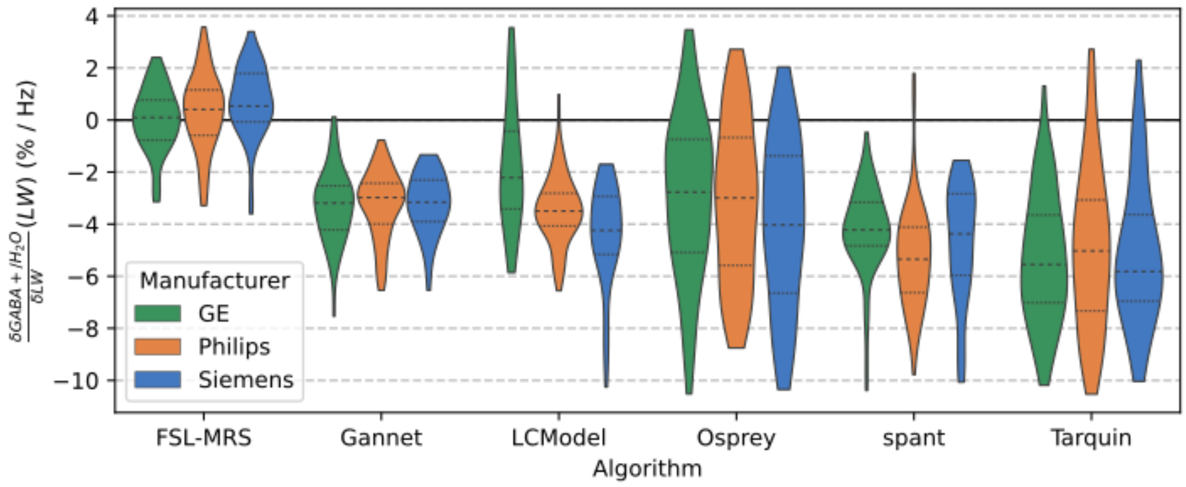


*Supplementary Figure 6 LLWF (rate of change in GABA+ estimate in relation to Lorentzian linewidth) for each subject, grouped by algorithm and scanner manufacturer.*

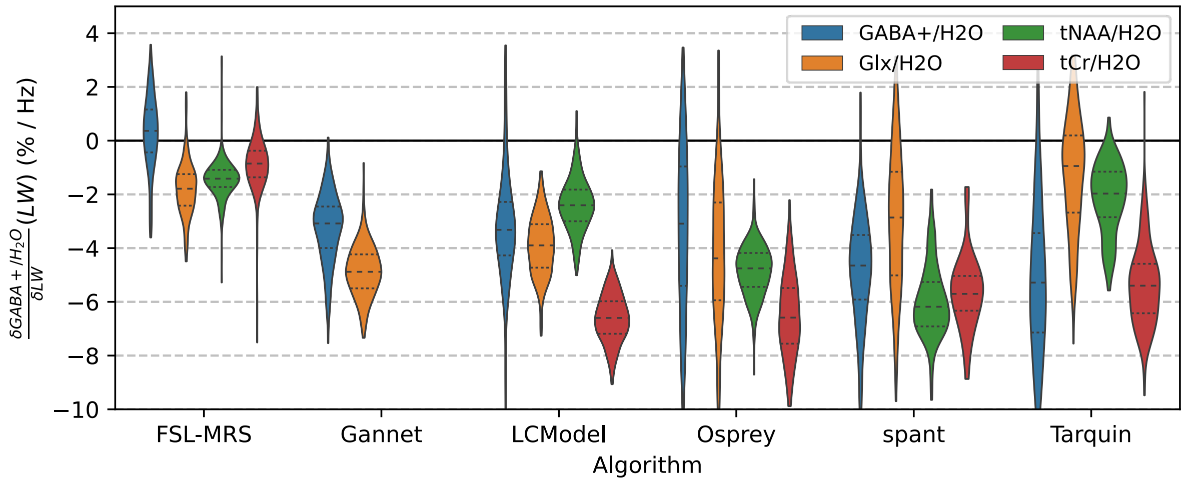


*Supplementary Figure 7 LLWF (rate of change in concentration estimate in relation to Lorentzian linewidth) assessed for alternative metabolites*

| Algorithm 1 | LLWF 1 | Algorithm 2 | LLWF 2 | tstat | p | p_holm | sig |
|---|---|---|---|---|---|---|---|
| FSL-MRS | 0.39 +- 1.4 %/Hz | Gannet | -3.25 +- 1.3 %/Hz | 26,38 | 8,987E-91 | 1,258E-89 | *** |
| FSL-MRS | 0.39 +- 1.4 %/Hz | LCModel | -3.16 +- 2.1 %/Hz | 19,7 | 6,193E-60 | 7,432E-59 | *** |
| FSL-MRS | 0.39 +- 1.4 %/Hz | Osprey | -3.05 +- 3.2 %/Hz | 13,5 | 3,868E-32 | 4,254E-31 | *** |
| FSL-MRS | 0.39 +- 1.4 %/Hz | Tarquin | -5.09 +- 2.9 %/Hz | 24,16 | 1,927E-72 | 2,505E-71 | *** |
| FSL-MRS | 0.39 +- 1.4 %/Hz | spant | -4.83 +- 2.0 %/Hz | 30,29 | 2,505E-104 | 3,757E-103 | *** |
| Gannet | -3.09 +- 1.3 %/Hz | LCModel | -3.16 +- 2.1 %/Hz | -0,517 | 0,6055 | 1 | |
| Gannet | -3.09 +- 1.3 %/Hz | Osprey | -3.05 +- 3.2 %/Hz | -0,834 | 0,4051 | 1 | |
| Gannet | -3.09 +- 1.3 %/Hz | Tarquin | -5.09 +- 2.9 %/Hz | 8,402 | 2,12E-15 | 1,696E-14 | *** |
| Gannet | -3.09 +- 1.3 %/Hz | spant | -4.83 +- 2.0 %/Hz | 9,652 | 8,402E-20 | 8,402E-19 | *** |
| LCModel | -3.32 +- 2.1 %/Hz | Osprey | -3.05 +- 3.2 %/Hz | -0,4294 | 0,6679 | 1 | |
| LCModel | -3.32 +- 2.1 %/Hz | Tarquin | -5.09 +- 2.9 %/Hz | 7,823 | 5,296E-14 | 3,707E-13 | *** |
| LCModel | -3.32 +- 2.1 %/Hz | spant | -4.83 +- 2.0 %/Hz | 8,386 | 7,514E-16 | 6,763E-15 | *** |
| Osprey | -2.99 +- 3.2 %/Hz | Tarquin | -5.09 +- 2.9 %/Hz | 6,761 | 4,97E-11 | 2,982E-10 | *** |
| Osprey | -2.99 +- 3.2 %/Hz | spant | -4.83 +- 2.0 %/Hz | 6,735 | 7,427E-11 | 3,714E-10 | *** |
| Tarquin | -5.26 +- 2.9 %/Hz | spant | -4.83 +- 2.0 %/Hz | -1,07 | 0,2856 | 1 | |

*Supplementary Table 5 Groupwise statistics for GABA+/$H_2O$ LLWF (see section 3.1.2)*

| Algorithm | Vendor 1 | LLWF 1 | Vendor 2 | LLWF 2 | tstat | p | p_holm | sig |
|---|---|---|---|---|---|---|---|---|
| FSL-MRS | G | 0.09 +- 1.3 %/Hz | P | 0.30 +- 1.5 %/Hz | -1,346 | 0,1804 | 1 | |
| FSL-MRS | G | 0.09 +- 1.3 %/Hz | S | 0.75 +- 1.3 %/Hz | -3,028 | 0,003123 | 0,0468 | * |
| FSL-MRS | P | 0.52 +- 1.5 %/Hz | S | 0.75 +- 1.3 %/Hz | -1,817 | 0,07202 | 0,8642 | |
| Gannet | G | -3.19 +- 1.4 %/Hz | P | -3.25 +- 1.3 %/Hz | -0,03473 | 0,9724 | 1 | |
| Gannet | G | -3.19 +- 1.4 %/Hz | S | -3.23 +- 1.2 %/Hz | -0,1225 | 0,9028 | 1 | |
| Gannet | P | -2.98 +- 1.3 %/Hz | S | -3.23 +- 1.2 %/Hz | -0,1046 | 0,9168 | 1 | |
| LCModel | G | -2.38 +- 2.4 %/Hz | P | -3.45 +- 1.3 %/Hz | 4,85 | 5.323E-06 | 9E-05 | *** |
| LCModel | G | -2.38 +- 2.4 %/Hz | S | -4.27 +- 2.1 %/Hz | 5,674 | 1,113E-07 | 2E-06 | *** |
| LCModel | P | -3.50 +- 1.3 %/Hz | S | -4.27 +- 2.1 %/Hz | 2,491 | 0,01512 | 0,1965 | |
| Osprey | G | -1.90 +- 3.0 %/Hz | P | -2.96 +- 3.2 %/Hz | 1,066 | 0,2884 | 1 | |
| Osprey | G | -1.90 +- 3.0 %/Hz | S | -3.99 +- 3.3 %/Hz | 2,537 | 0,01281 | 0,1793 | |
| Osprey | P | -3.02 +- 3.2 %/Hz | S | -3.99 +- 3.3 %/Hz | 1,767 | 0,0807 | 0,8877 | |
| Tarquin | G | -5.57 +- 2.4 %/Hz | P | -4.92 +- 3.1 %/Hz | -0,9852 | 0,3261 | 1 | |
| Tarquin | G | -5.57 +- 2.4 %/Hz | S | -5.08 +- 2.8 %/Hz | -0,5701 | 0,5699 | 1 | |
| Tarquin | P | -5.01 +- 3.1 %/Hz | S | -5.08 +- 2.8 %/Hz | 0,3065 | 0,7598 | 1 | |
| spant | G | -4.25 +- 1.7 %/Hz | P | -5.26 +- 2.0 %/Hz | 3,24 | 0,001474 | 0,0236 | * |
| spant | G | -4.25 +- 1.7 %/Hz | S | -4.67 +- 2.2 %/Hz | 0,9545 | 0,3424 | 1 | |
| spant | P | -5.35 +- 2.0 %/Hz | S | -4.67 +- 2.2 %/Hz | -1,576 | 0,1185 | 1 | |

*Supplementary Table 6 Groupwise statistics for GABA+/$H_2O$ LLWF, compared within algorithm across sites*

### D Supplementary Details: Synthetic Data

#### D.1 Synthetic data generation

Further to details in the main manuscript, section 2.3: Background signal derived by this approach is only meaningful on the original fit range (0.2-4.2 ppm); this has implications for the measurement of SNR, which often uses a region well beyond the normal fit range to assess the noise factor. Moreover, there is a risk of discontinuities (hence potential numeric instability) if the background function is not close to zero at the edge of the fit range. To avoid such discontinuities, a smooth spline was fit to the last 0.05 ppm at either end of the fit range, and extrapolated 0.10 ppm beyond the fit range. A pair of Hanning functions ($\cos^2 \frac{\pi(f-f_0)}{2a}$) were used to transition first from the signal to the smooth spline fit in the last 0.05 ppm, and then from the smooth spline fit to zero in the subsequent 0.10 ppm beyond the fit range.

#### D.2 Outcomes from fits to synthetic data

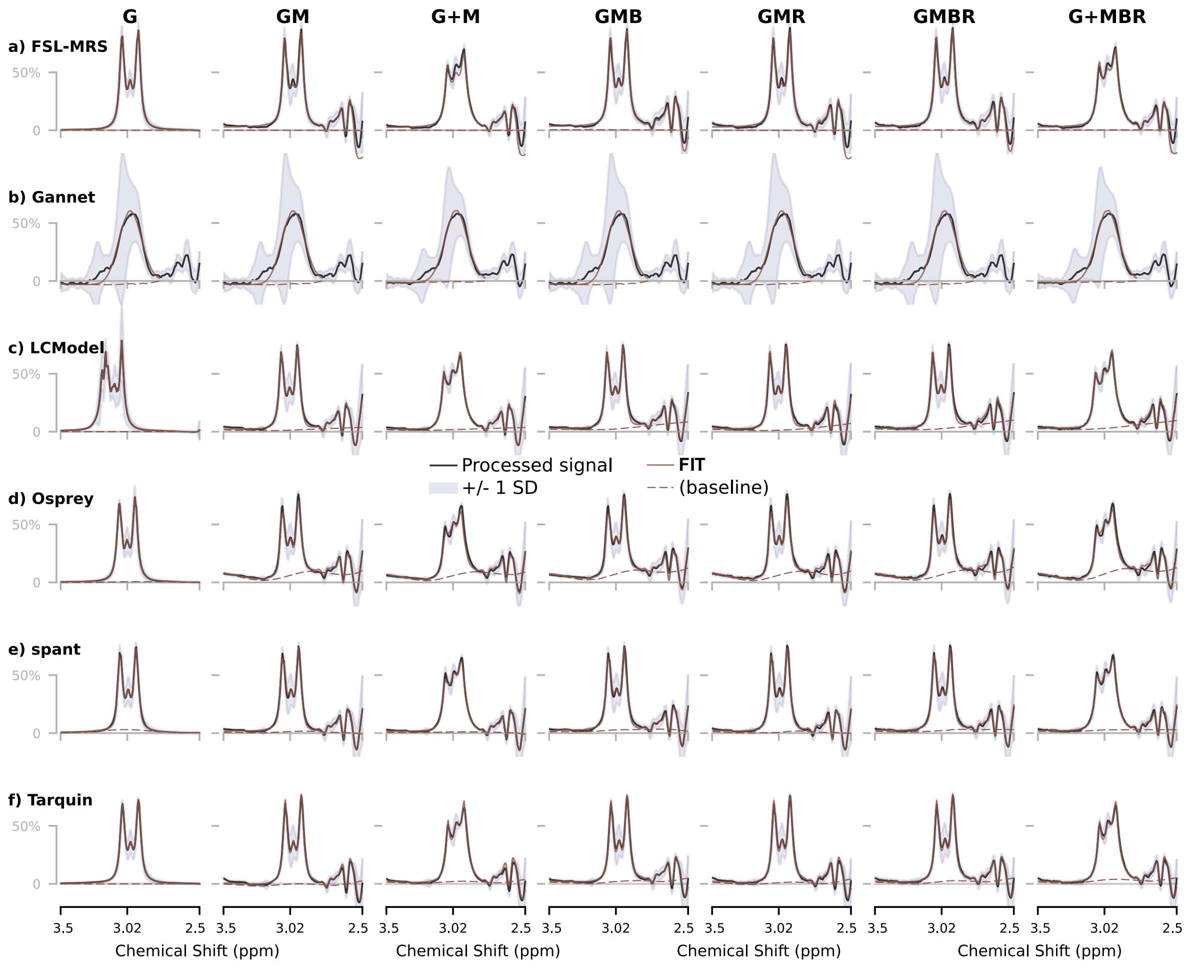


*Supplementary Figure 8 Synthetic spectra and fits by various algorithms*

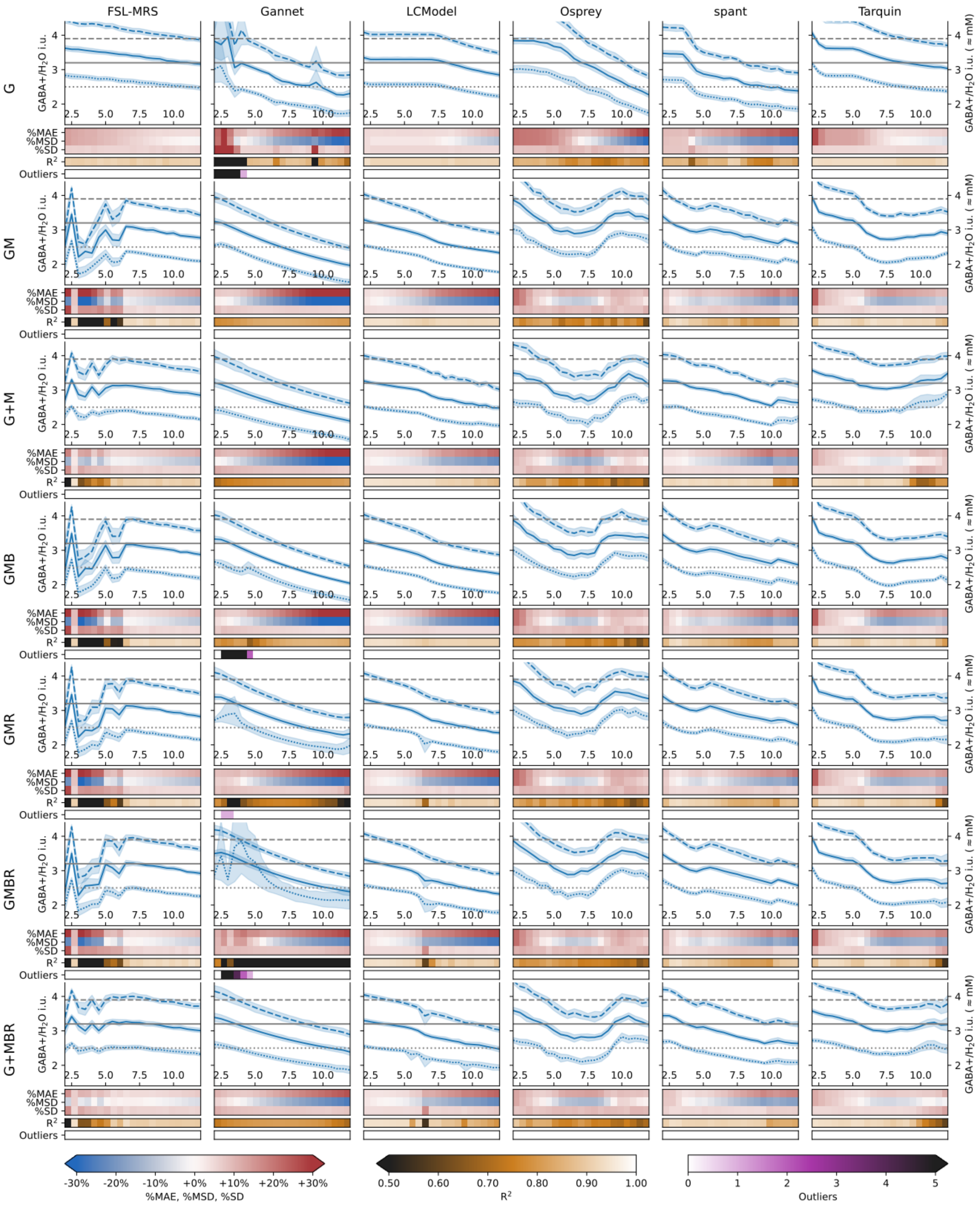


*Supplementary Figure 9 GABA+/H2O estimates from each algorithm as a function of Lorentzian linewidth, based on synthetic data models of varying complexities, for nominal GABA+ concentrations as detailed in Section 2.3.*

### E Exploratory Analysis #1: spant baseline modelling

#### E.1 Background and Methods

The default fitting algorithm in spant is ABfit. A key component of this algorithm is the adaptive baseline model, in which the degree of baseline flexibility is automatically adjusted to match the complexity of the data[1]. As with other algorithms, this is inevitably a balancing act between a baseline that is too smooth, and one that is too flexible – with either scenario potentially leading to bias or increased variance in the fit outcome. Most pre-existing literature and documentation relating to spant at the time of this analysis focussed on non-edited spectra, such as may be obtained by short TE PRESS sequence. Early piloting suggested that default parameters suitable for such sequences may have allowed for too much baseline flexibility for GABA-edited datasets – as such, several different configurations of spant were applied to the in-vivo datasets to assess their relative suitability for GABA-edited spectra.

There are a number of options available for controlling ABfit's baseline modelling; two of these are `aic_smoothing_factor` and `bl_ed_ppm`:

- `aic_smoothing_factor` affects the criterion used for automatic estimation, with smaller values encouraging more flexible baselines, while remaining adaptive to any broad spectral features. Five different options for `aic_smoothing_factor` were assessed: 5 (default), 10, 20, 40 and 80, specifying increasingly firm baselines; these are subsequently denoted A05, A10, A20, A40 and A80 respectively.

- `bl_ed_pppm` specifies the baseline flexibility, in units of ED per ppm where ED is the "effective dimension" – analogous to the number of spline functions required per ppm; this is automatically determined except when `auto_bl_flex` (automatic baseline flexibility) is explicitly disabled. Three fixed values for `bl_ed_pppm` were tested (with `auto_bl_flex` disabled): 1.00, 1.67 and 2.00 ED/ppm, denoted F10, F17 and F20.

[1] These parameters are detailed at https://martin3141.github.io/spant/articles/abfit-baseline-opts.html

This analysis was performed using all in-vivo datasets from the main study, with processing and quality control as described in the main manuscript – including spectra with Lorentzian linebroadening, but not including those of experimentally degraded SNR.

To assess the relative suitability of each configuration, the determined baseline flexibility (`bl_ed_pppm`) and resultant signal-to-residual ratio (SRR) were inspected. Furthermore, GABA+ estimates were assessed for accuracy and consistency with other algorithms methods[2]. Exploiting the different expected GABA concentrations between grey and white matter[3–5], the degree of correlation between GABA estimates and voxel tissue fraction was used as an index of the accuracy of GABA estimation[6]. Robust (skipped)[7,8] Spearman correlation coefficients between voxel grey matter fraction and water-scaled GABA estimate without adjustment for tissue content (GABA+/$H2O_{noTC}$) were reported. Finally, intra-class correlation coefficients (ICCs) were calculated between each spant variant and the median outcome from all other algorithms, using a two-way mixed-effects model for single-rater consistency (ICC(3,1), implemented in pingouin[9]).

#### E.2 Results and Discussion

Outcomes for each of the tested configurations are presented in Supplementary Table 7. Using automatic baseline flexibility (A*nn*), the determined `bl_ed_pppm` factor was seen to clip at either edge of the allowable range (.502, 7.00) in a substantial proportion of cases; with A05 case (`aic_smoothing_factor`=5), the majority of estimates clipped to the predefined upper limit; in the A80 case, the majority clipped to the lower limit. The reported GABA.sd metric is suspiciously low in several cases (particularly with lower `bl_ed_pppm` values); as such, this metric is not considered further herein, but may warrant further investigation elsewhere. Signal-to-residual ratio (SRR) was generally higher in the cases with greater baseline flexibility, in particular A05. This is to be expected: more flexible baselines are more able to absorb signal fluctuations which would otherwise remain in the residuals. The intraclass correlation coefficient (ICC), indicating agreement with other algorithms, was highest in the fixed `bl_ed_pppm` cases (F*nn*), particularly F17 (`bl_ed_pppm`=1.67); this configuration is roughly equivalent to the 0.6 ppm baseline knot spacing adopted in both LCModel and Osprey, hence it is unsurprising that greatest agreement is seen in this case, where the baseline parameterisation is most similar. Using correlation between

GABA+/H2O$_{noTC}$ and tissue grey matter (GMcorr) as an indication of measurement accuracy, although subtle variation was seen, none of the configurations was found to be significantly more accurate than any other. The impact of line broadening, as assessed with the LLWF (see main manuscript section 2.5), was substantially reduced in magnitude as baseline flexibility was reduced.

| Subset | parameter | A05 | A10 | A20 | A40 | A80 | F10 | F17 | F20 |
|---|---|---|---|---|---|---|---|---|---|
| Unfiltered only | GABA+/H2O | 3.39 [3.02; 3.81] | 3.42 [3.06; 3.84] | 3.78 [3.33; 4.16] | 3.87 [3.47; 4.42] | 3.91 [3.48; 4.38] | 3.86 [3.53; 4.37] | 3.69 [3.42; 4.08] | 3.56 [3.29; 3.95] |
| | aic_smoothing_factor | 5 | 10 | 20 | 40 | 80 | *N/A* | *N/A* | *N/A* |
| | bl_ed_pppm | 7.00 [7.00; 7.00] | 7.00 [4.09; 7.00] | 1.35 [0.502; 5.89] | 0.502 [0.502; 1.93] | 0.502 [0.502; 0.503] | *1* | *1,67* | *2* |
| | SRR | 26.8 [13.1; 148.] | 17.1 [10.6; 23.4] | 26.4 [9.98; 112.] | 11.0 [8.15; 14.7] | 10.5 [7.60; 14.0] | 10.8 [7.52; 14.4] | 11.7 [8.10; 15.5] | 12.1 [8.45; 16.1] |
| | GABA.sd | 1.35 [0.411; 4.45] | 0.622 [0.350; 1.71] | 0.0101 [6.62e-09; 1.27] | 1.78e-08 [1.24e-8; 0.235] | 1.75e-08 [1.2e-8; 2.7e-8] | 2.33e-08 [1.39e-8; 0.001] | 0.188 [0.0121; 0.266] | 0.250 [0.178; 0.853] |
| | Gmcorr | 0,364 | 0,396 | 0,346 | 0,377 | 0,4 | 0,363 | 0,368 | 0,372 |
| | ICC | 0.407 {0.290; 0.510} | 0.438 {0.320; 0.540} | 0.580 {0.480; 0.660} | 0.641 {0.560; 0.710} | 0.636 {0.550; 0.710} | 0.711 {0.640; 0.770} | 0.675 {0.600; 0.740} | 0.665 {0.580; 0.730} |
| All (filtered and unfiltered) | GABA+/H2O | 2.95 [2.25; 3.47] | 2.98 [2.37; 3.50] | 3.13 [2.57; 3.73] | 3.49 [2.77; 4.11] | 3.65 [2.99; 4.25] | 3.64 [3.02; 4.32] | 3.41 [2.95; 3.89] | 3.26 [2.78; 3.74] |
| | bl_ed_pppm | 7.00 [7.00; 7.00] | 7.00 [5.89; 7.00] | 4.92 [0.825; 7.00] | 0.626 [0.502; 3.39] | 0.502 [0.502; 1.61] | *1* | *1,67* | *2* |
| | SRR | 44.7 [19.6; 198.] | 24.3 [16.4; 35.4] | 44.0 [15.0; 161.] | 12.3 [9.14; 16.7] | 10.2 [7.07; 13.8] | 10.5 [6.23; 14.3] | 12.1 [7.96; 15.9] | 12.9 [8.84; 16.9] |
| | GABA.sd | 0.740 [0.252; 2.55] | 0.395 [0.215; 1.28] | 0.478 [5.98e-05; 1.87] | 2.38e-08 [8.62e-9; 0.381] | 1.32e-08 [7.09e-9; 0.004] | 1.72e-08 [7.81e-9; 0.0004] | 0.108 [0.00415; 0.198] | 0.167 [0.0878; 0.603] |
| | LLWF | -5.78 {-6.02; -5.52} | -5.69 {-5.95; -5.40} | -4.54 {-4.75; -4.32} | -3.00 {-3.27; -2.80} | -2.85 {-3.03; -2.53} | -3.10 {-3.29; -2.83} | -3.99 {-4.34; -3.75} | -4.35 {-4.54; -4.06} |

*Supplementary Table 7 Selected fitting performance metrics for spant, using a variety of baseline configurations. Values are reported as median with [10,90]-percentiles or {95% confidence interval}.*

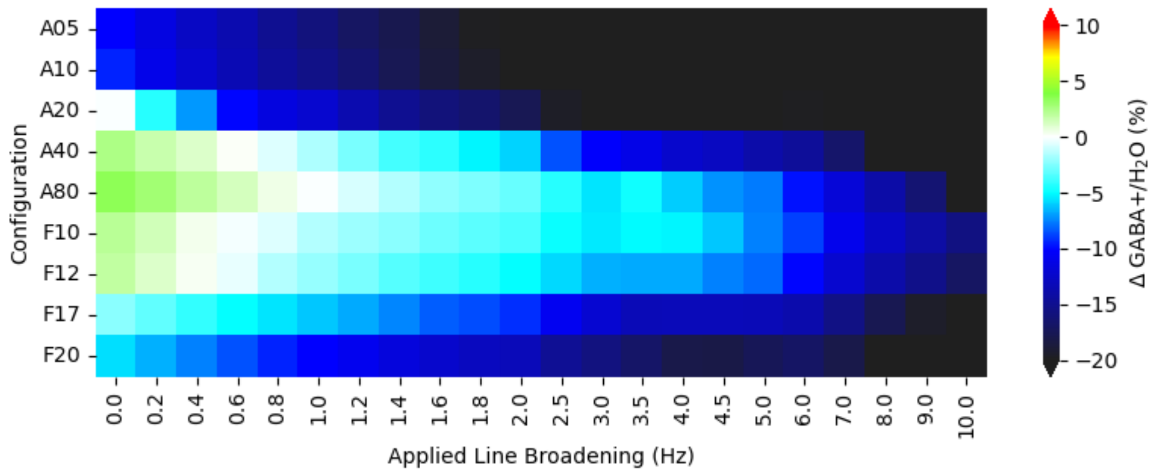


*Supplementary Figure 10 Relative difference in GABA+ concentration reported by spant in different configurations, in relation to applied line broadening*

Although certain configurations displayed better performance for individual metrics, or for individual sites' datasets, no single configuration was found to clearly outperform the others across a range of quality metrics in the general case. For settings where linewidth differences are not expected to be an issue, the default configuration (A05) remains a reasonable choice; in cases where linewidth changes may be of concern, a more firm baseline model should be adopted.

In the present study, configuration A20 was adopted (`aic_smoothing_factor`=20) for the main analysis: this configuration is seen to allow the Abfit algorithm to operate over its configured range (rather than clipping significantly at either limit), has a 'typical' baseline resolution similar to that of other methods, and gives moderately good outcomes across a range of quality metrics without optimising for any particular factor.

### F Exploratory Analysis #2: FSL-MRS baseline modelling, optimization and lineshape

#### F.1 Background and Methods

FSL-MRS exhibited two noteworthy characteristics in the main analysis:

- For the filtered in-vivo datasets, fit outcomes showed notably lower linewidth dependency (for Lorentzian linebroadening) than for any of the other configurations assessed.
- Somewhat greater variance was seen for datasets of extremely low linewidth (particularly apparent in the simulated datasets)

To investigate the first and mitigate the second, several alternative configurations of FSL-MRS were assessed:

- Polynomial baseline order (`--baseline_order`) of -1 (no baseline), 1 (linear) and 2-7 (*n*th order polynomials) were assessed
- Different optimization algorithms (`--algo`) were evaluated: Truncated Newton (“Newton”) and full Metropolis Hastings (“MH”). In typical usage, the latter is initialised internally by the substantially faster Newton algorithm.
- Modelling without and with a separate metabolite group for NAA and NAAG (`--metab_groups NAA+NAAG`), to allow shift and broadening of those metabolites to be optimised separately from all other components (hence allowing a slight frequency shift to mitigate the observed instability at very low linewidths)
- Comparing the `--lorentzian` lineshape model against the default Voigt lineshape model: the former may be a closer match to the line broadening applied in the present analysis, whereas the latter is expected to more fully reflect the different broadening factors present in in-vivo data.

As with the spant baseline exploration, the same in-vivo datasets, processing and quality control as described in the main manuscript were used – including spectra with Lorentzian linebroadening spectra, but not including those of experimentally degraded SNR.

#### F.2 Results and Discussion

Lower-order baseline models (below 5$^{th}$ order) exhibit the lowest linewidth dependence for moderate line-broadening factors, up to about 5 Hz; slightly increased linewidth dependence was seen with higher-order baseline models. With higher order baselines ascribing progressively more signal to baseline, the default 2$^{nd}$ order baseline remains a reasonable choice for scenarios where moderate linewidth differences may be encountered. Adopting a pure Lorentzian lineshape model resulted in only marginal differences, with a tendency towards fractionally higher estimates and perhaps a slightly reduced linewidth dependence for moderate linewidths, in cases where a baseline was modelled (not statistically significant).

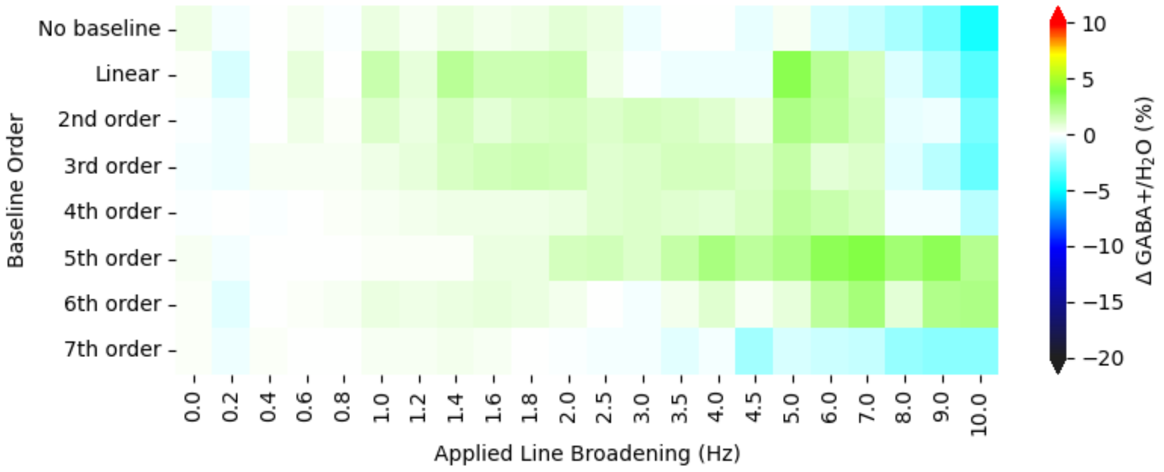


*Supplementary Figure 11 Relative difference in GABA+ concentration reported by FSL-MRS with differing baseline configurations and default Lorentzian + Gaussian linewidth model, in relation to applied line broadening*

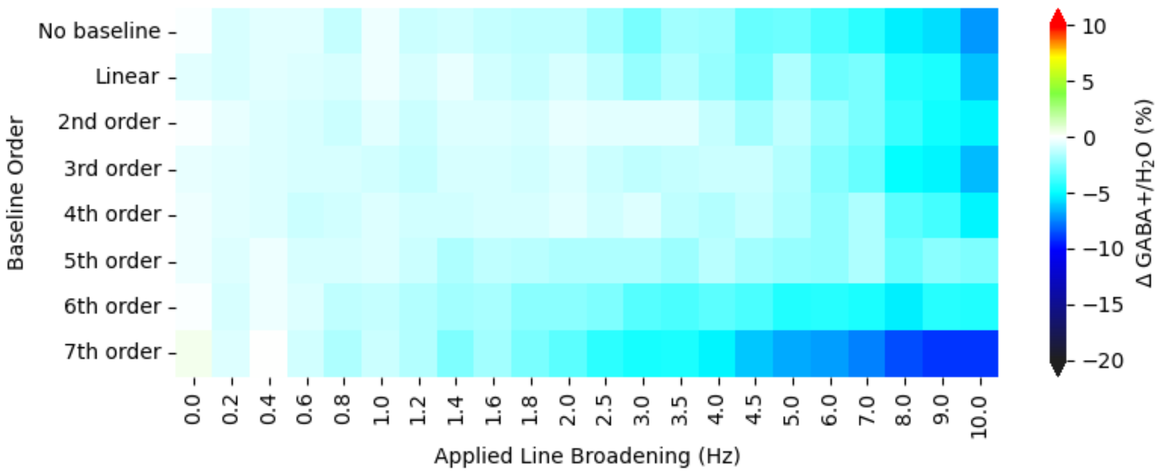


*Supplementary Figure 12 Relative difference in GABA+ concentration reported by FSL-MRS with differing baseline configurations and pure Lorentzian lineshape model, in relation to applied line broadening*

The GABA+ model sub-parts (GABA and MM3co) exhibit complementary behaviours which combine to give a relatively flat GABA+ response; estimates for these constituent parts are more stable (in relation to linewidth) when a 2$^{nd}$ order baseline is modelled. However, at lower linewidths (<2 Hz broadening) some additional variance in the estimates appears, particularly in cases where a baseline is modelled. This is reduced (but not entirely eliminated) when using the full MH model, which additionally exhibits slightly reduced linewidth dependence.

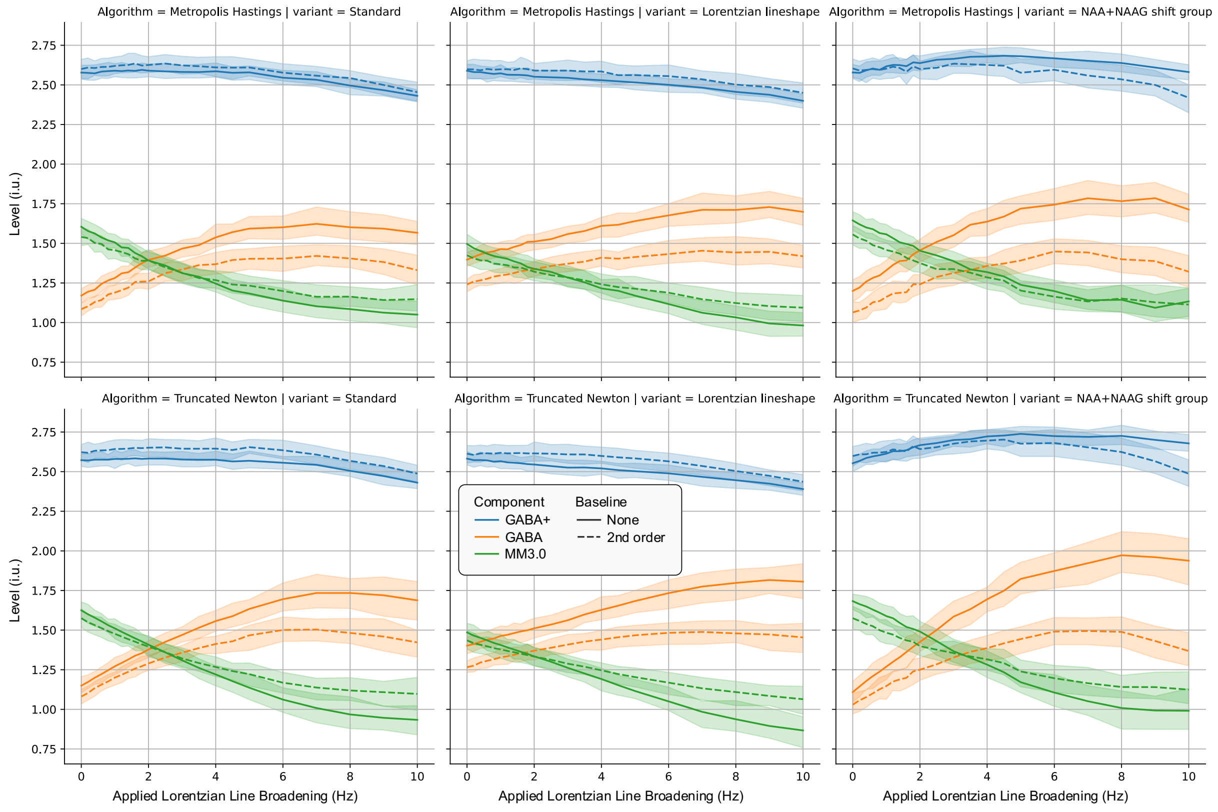


*Supplementary Figure 13 Relative GABA+ and partial GABA/MM3co estimates from FSL-MRS under varying linewidth models (Voigt or Lorentzian), differing baseline configurations (2nd order or None), and differing optimization strategies (truncated Newton or full Metropolis-Hastings)*

| Baseline Model | algo | Variant | Unfiltered only GABA+/H2O | GMcorr | ICC | All (filtered and unfiltered) GABA+/H2O | LLWF |
|---|---|---|---|---|---|---|---|
| No baseline | Newton | | 2.60 [2.16; 2.92] | 0,356 | 0.754 {0.690; 0.810} | 2.58 [2.12; 2.97] | 0.188 {0.0755; 0.349} |
| Linear | Newton | | 2.59 [2.14; 3.03] | 0,306 | 0.854 {0.810; 0.890} | 2.60 [2.11; 3.08] | 0.396 {0.309; 0.630} |
| 2nd order | Newton | | 2.58 [2.14; 3.14] | 0,251 | 0.860 {0.820; 0.890} | 2.61 [2.10; 3.18] | 0.407 {0.184; 0.602} |
| 3rd order | Newton | | 2.57 [2.10; 3.16] | 0,296 | 0.844 {0.800; 0.880} | 2.60 [2.04; 3.20] | 0.217 {0.0221; 0.472} |
| 4th order | Newton | | 2.58 [2.07; 3.12] | 0,327 | 0.833 {0.790; 0.870} | 2.59 [2.04; 3.19] | 0.376 {0.110; 0.470} |
| 5th order | Newton | | 2.59 [2.12; 3.12] | 0,313 | 0.775 {0.720; 0.820} | 2.61 [2.09; 3.19] | 0.309 {0.0703; 0.482} |
| 6th order | Newton | | 2.58 [2.06; 3.09] | 0,389 | 0.704 {0.630; 0.770} | 2.60 [2.06; 3.15] | 0.189 {-0.0378; 0.425} |
| 7th order | Newton | | 2.59 [2.05; 3.11] | 0,413 | 0.695 {0.620; 0.760} | 2.57 [2.03; 3.12] | **-0.0392 {-0.245; 0.0781}** |
| No baseline | Newton | NAA+NAAG | 2.56 [2.15; 3.00] | 0,432 | 0.719 {0.650; 0.780} | 2.71 [2.22; 3.28] | 2.42 {2.05; 2.76} |
| Linear | Newton | NAA+NAAG | 2.53 [2.10; 3.10] | 0,294 | 0.807 {0.750; 0.850} | 2.71 [2.12; 3.44] | 2.35 {1.91; 2.86} |
| 2nd order | Newton | NAA+NAAG | 2.52 [2.09; 3.27] | 0,271 | 0.843 {0.800; 0.880} | 2.65 [2.05; 3.47] | 2.04 {1.70; 2.56} |
| 3rd order | Newton | NAA+NAAG | 2.54 [2.05; 3.32] | **0,237** | 0.860 {0.820; 0.890} | 2.66 [2.07; 3.49] | 2.57 {2.07; 2.84} |
| 4th order | Newton | NAA+NAAG | 2.56 [2.04; 3.26] | 0,281 | 0.829 {0.780; 0.870} | 2.67 [2.11; 3.50] | 2.90 {2.37; 3.29} |
| 5th order | Newton | NAA+NAAG | 2.55 [2.03; 3.18] | 0,278 | 0.812 {0.760; 0.850} | 2.67 [2.11; 3.49] | **2.94 {2.23; 3.37}** |
| 6th order | Newton | NAA+NAAG | 2.55 [2.02; 3.21] | 0,364 | 0.735 {0.670; 0.790} | 2.71 [2.10; 3.48] | 2.46 {2.01; 3.00} |
| 7th order | Newton | NAA+NAAG | 2.55 [2.06; 3.25] | 0,376 | **0.665 {0.580; 0.730}** | 2.71 [2.08; 3.47] | 1.92 {1.54; 2.45} |
| No baseline | Newton | Lorentzian | 2.60 [2.17; 2.96] | 0,309 | 0.715 {0.640; 0.770} | 2.55 [2.10; 2.93] | -0.728 {-0.874; -0.574} |
| Linear | Newton | Lorentzian | 2.59 [2.14; 3.02] | 0,304 | 0.849 {0.810; 0.880} | 2.57 [2.08; 3.02] | -0.250 {-0.419; -0.159} |
| 2nd order | Newton | Lorentzian | 2.60 [2.14; 3.11] | 0,273 | **0.869 {0.830; 0.900}** | 2.57 [2.07; 3.12] | -0.174 {-0.306; 0.103} |
| 3rd order | Newton | Lorentzian | 2.59 [2.09; 3.18] | 0,266 | 0.858 {0.820; 0.890} | 2.56 [2.01; 3.18] | -0.169 {-0.344; 0.0912} |
| 4th order | Newton | Lorentzian | 2.59 [2.08; 3.14] | 0,336 | 0.839 {0.790; 0.870} | 2.57 [2.01; 3.17] | -0.259 {-0.487; -0.120} |
| 5th order | Newton | Lorentzian | 2.59 [2.12; 3.06] | 0,325 | 0.799 {0.750; 0.840} | 2.56 [2.02; 3.11] | -0.597 {-0.826; -0.420} |
| 6th order | Newton | Lorentzian | 2.60 [2.13; 3.05] | 0,423 | 0.729 {0.660; 0.790} | 2.54 [2.01; 3.04] | -1.08 {-1.27; -0.862} |
| 7th order | Newton | Lorentzian | 2.61 [2.11; 3.07] | 0,435 | 0.732 {0.660; 0.790} | 2.50 [1.95; 2.99] | -1.59 {-1.80; -1.39} |
| No baseline | MH | | 2.60 [2.15; 2.96] | 0,323 | 0.807 {0.750; 0.850} | 2.58 [2.12; 2.98] | 0.220 {0.00176; 0.399} |
| 2nd order | MH | | 2.56 [2.09; 3.15] | 0,259 | 0.834 {0.790; 0.870} | 2.58 [2.03; 3.18] | 0.276 {0.118; 0.527} |
| No baseline | MH | NAA+NAAG | 2.55 [2.18; 3.04] | 0,414 | 0.764 {0.700; 0.810} | 2.67 [2.18; 3.25] | 1.58 {1.27; 1.95} |
| 2nd order | MH | NAA+NAAG | 2.57 [2.01; 3.22] | 0,245 | 0.833 {0.790; 0.870} | 2.58 [1.98; 3.44] | 1.23 {0.741; 1.78} |
| No baseline | MH | Lorentzian | 2.60 [2.20; 3.01] | 0,368 | 0.719 {0.650; 0.780} | 2.55 [2.10; 2.95] | -0.526 {-0.679; -0.391} |
| 2nd order | MH | Lorentzian | 2.58 [2.06; 3.15] | 0,254 | 0.831 {0.780; 0.870} | 2.56 [2.01; 3.15] | -0.0440 {-0.356; 0.168} |

*Supplementary Table 8 Quality metrics for FSL-MRS modelling outcomes with differing baseline and algorithm options*

### G Exploratory Analysis #3: LCModel baseline knot spacing

#### G.1 Background and Methods

To assess the impact of baseline flexibility on linewidth dependence in LCModel, spectra were fit using a range of different baseline configurations: no baseline (`NOBASE=T`), and spline knot spacings from 0.1 to 4.0 ppm (in steps of 0.1 ppm between 0.1 to 1.2 ppm, steps of 0.2 ppm between 1.2 and 2.0 ppm, and in steps of 0.5 ppm from 2.0 up to 4.0 ppm). As previously, in-vivo data sets were fit after processing as described in the main manuscript, including spectra with Lorentzian linebroadening but not including those of experimentally degraded SNR.

#### G.2 Results and Discussion

GABA+ estimates obtained when no baseline was modelled were significantly elevated relative to those obtained when also using a baseline model. Unsurprisingly, excessively flexible baseline models (< 0.4 ppm knot spacing) resulted in significantly stronger linewidth-dependent biases. Firmer baseline models (>0.4 ppm) led to lower linewidth-dependent bias, particularly in the 0.5 – 1.0 ppm knot spacing range; this is compatible with previous recommendations for knot spacing[2,10] in the order of 0.55-0.6 ppm.

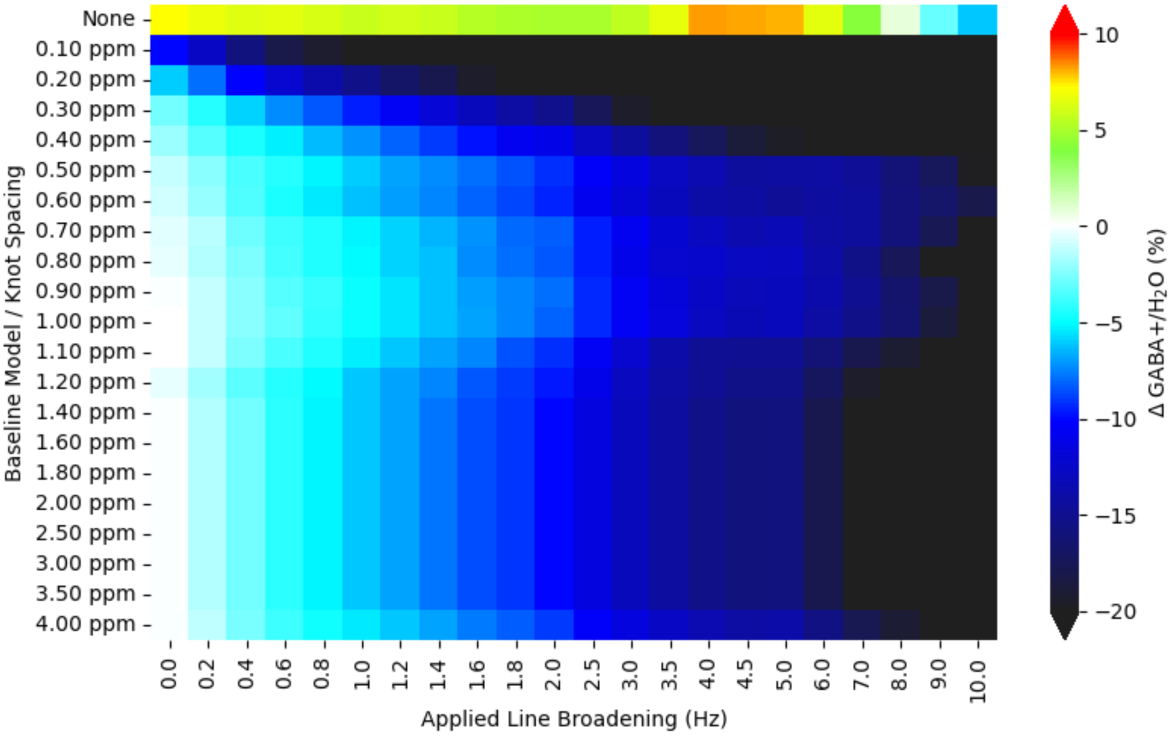


*Supplementary Figure 14 Relative difference in GABA+ concentration reported by LCModel with different baseline knot spacings, in relation to applied line broadening*

| Baseline Knot Spacing | Unfiltered only | | | | All (filtered and unfiltered) | | |
|---|---|---|---|---|---|---|---|
| | GABA+/H2O | GABA+ %SD | GMcorr | ICC | GABA+/H2O | GABA+ %SD | LLWF |
| None | 3.52 [3.16; 3.89] | 10.5 [7.52; 16.9] | **0,305** | 0.630 {0.540; 0.700} | 3.45 [2.78; 4.17] | 10.5 [4.00; 21.1] | **-1.16 {-1.47; -0.787}** |
| 0.10 ppm | 2.68 [2.36; 3.05] | 13.6 [10.3; 20.8] | 0,352 | **0.432 {0.320; 0.530}** | 2.34 [1.33; 3.22] | **12.6 [6.01; 25.6]** | -6.75 {-7.28; -6.05} |
| 0.20 ppm | 2.87 [2.57; 3.18] | 11.7 [8.64; 17.5] | 0,321 | 0.578 {0.480; 0.660} | 2.52 [1.60; 3.34] | 10.4 [5.78; 21.4] | **-7.61 {-8.14; -7.03}** |
| 0.30 ppm | 3.10 [2.83; 3.42] | 10.9 [7.88; 16.4] | 0,332 | 0.675 {0.600; 0.740} | 2.71 [1.76; 3.44] | 9.95 [5.00; 20.6] | -6.71 {-7.38; -6.26} |
| 0.40 ppm | 3.15 [2.89; 3.48] | 10.7 [7.30; 16.1] | 0,334 | 0.705 {0.630; 0.770} | 2.89 [2.23; 3.53] | 9.67 [4.84; 20.6] | -4.62 {-5.06; -4.26} |
| 0.50 ppm | 3.19 [2.92; 3.53] | 10.5 [7.00; 16.2] | 0,32 | 0.744 {0.680; 0.800} | 2.97 [2.36; 3.59] | **9.64 [4.59; 21.0]** | -3.76 {-3.96; -3.52} |
| 0.60 ppm | 3.19 [2.92; 3.53] | 10.6 [7.10; 16.0] | 0,35 | 0.745 {0.680; 0.800} | 2.97 [2.36; 3.60] | 9.89 [4.78; 21.0] | -4.02 {-4.31; -3.70} |
| 0.70 ppm | 3.20 [2.94; 3.55] | 10.5 [7.02; 15.6] | 0,33 | 0.731 {0.660; 0.790} | 2.99 [2.37; 3.62] | 9.77 [4.82; 20.9] | **-3.63 {-4.10; -3.24}** |
| 0.80 ppm | 3.22 [2.92; 3.56] | 10.5 [7.03; 15.8] | 0,342 | 0.745 {0.680; 0.800} | 2.98 [2.34; 3.61] | 9.76 [4.65; 20.9] | -3.90 {-4.18; -3.52} |
| 0.90 ppm | 3.22 [2.92; 3.55] | 10.4 [7.09; 15.7] | 0,352 | **0.747 {0.680; 0.800}** | 3.00 [2.37; 3.62] | 9.69 [4.50; 21.0] | -3.89 {-4.13; -3.40} |
| 1.00 ppm | 3.22 [2.91; 3.55] | 10.4 [7.10; 16.0] | 0,328 | 0.745 {0.680; 0.800} | 2.99 [2.36; 3.62] | 9.67 [4.32; 20.8] | -3.90 {-4.09; -3.47} |
| 1.10 ppm | 3.24 [2.84; 3.59] | 10.4 [7.09; 16.5] | 0,343 | 0.743 {0.680; 0.800} | 2.96 [2.20; 3.67] | 9.68 [4.09; 20.8] | -4.39 {-4.85; -3.97} |
| 1.20 ppm | 3.22 [2.85; 3.52] | 10.6 [7.04; 16.7] | 0,342 | 0.741 {0.670; 0.800} | 2.93 [2.21; 3.62] | 9.95 [4.26; 21.0] | -4.57 {-4.79; -4.08} |
| 1.40 ppm | 3.21 [2.86; 3.55] | 10.5 [7.09; 16.1] | **0,364** | 0.743 {0.680; 0.800} | 2.93 [2.16; 3.64] | 9.90 [4.00; 20.8] | -4.67 {-5.21; -3.97} |
| 1.60 ppm | 3.21 [2.86; 3.55] | 10.5 [7.09; 16.1] | **0,364** | 0.743 {0.680; 0.800} | 2.93 [2.16; 3.64] | 9.90 [4.00; 20.8] | -4.67 {-5.19; -3.98} |
| 1.80 ppm | 3.21 [2.86; 3.55] | 10.5 [7.09; 16.1] | **0,364** | 0.743 {0.680; 0.800} | 2.93 [2.16; 3.64] | 9.90 [4.00; 20.8] | -4.67 {-5.21; -3.98} |
| 2.00 ppm | 3.21 [2.86; 3.55] | 10.5 [7.09; 16.1] | **0,364** | 0.743 {0.680; 0.800} | 2.93 [2.16; 3.64] | 9.90 [4.00; 20.8] | -4.67 {-5.20; -4.11} |
| 2.50 ppm | 3.21 [2.86; 3.55] | 10.5 [7.09; 16.1] | **0,364** | 0.743 {0.680; 0.800} | 2.93 [2.16; 3.64] | 9.90 [4.00; 20.8] | -4.67 {-5.19; -3.98} |
| 3.00 ppm | 3.21 [2.86; 3.55] | 10.5 [7.09; 16.1] | **0,364** | 0.743 {0.680; 0.800} | 2.93 [2.16; 3.64] | 9.90 [4.00; 20.8] | -4.67 {-5.19; -3.98} |
| 3.50 ppm | 3.21 [2.86; 3.55] | 10.5 [7.09; 16.1] | **0,364** | 0.743 {0.680; 0.800} | 2.93 [2.16; 3.64] | 9.90 [4.00; 20.8] | -4.67 {-5.19; -3.98} |
| 4.00 ppm | 3.21 [2.93; 3.53] | 10.5 [7.09; 15.8] | 0,348 | 0.730 {0.660; 0.790} | 2.96 [2.33; 3.59] | 9.85 [4.00; 20.4] | -4.12 {-4.44; -3.80} |

*Supplementary Table 9 Quality metrics for LCModel modelling outcomes with differing baseline configurations*

### H Supplementary Methods

Further details on specific parameters adopted for each of the modelling algorithms are presented below; details in this section have been published previously[2], and are reproduced here for completeness. This supplements detail provided in the main article, Section 2.4. Section H.1 focusses on modelling of the GABA-edited difference spectrum, with adaptions for the edit-OFF case mentioned briefly in Section H.2 where applicable.

#### H.1 Difference spectrum modelling

##### H.1.1 FSL-MRS

FSL-MRS [11] models data as a linear combination of basis components in the frequency domain, using Bayesian statistics for optimization. Version 1.1.1 was used, with the standard basis set as described in section 2.2 and the recommended Metropolis-Hastings algorithm on the default fit range (0.2-4.2 ppm).

Default baseline model is a second-order polynomial fit over the defined fit range. To allow FSL-MRS to yield water-scaled estimates, NAA+NAAG was defined as an internal reference for basis set scaling, and fixed tissue content (50% white/50% grey matter, 0% CSF) were supplied; this scaling was subsequently reversed, before re-scaling with per-subject tissue content. Alternative baseline configurations and variations to the “shift group” and algo parameters are explored in Supplementary Section F.

Typical invocation of FSL-MRS for GABA-edited difference spectra was with the following parameters:

```
--basis ge_diff_8192_4000.basis
--data S12_GABA_68.diff.nii.gz
--h2o S12_GABA_68.ref.nii.gz
--output S12_GABA_68.diff
--overwrite
--algo MH
--TE 68
--TR 2.0
--tissue_frac .5 .5 0
--internal_ref NAA NAAG
--verbose
--report
--combine NAA NAAG
--combine Glu Gln GSH
--combine GABA MM3co
```

```
Defaults:
  --ppmlim:           (0.2, 4.2)
  --baseline_order:   2
  --metab_groups:     0
  --h2o_scale:        1.0
```

##### H.1.2 Gannet

Gannet [12] models data by fitting peaks in the frequency domain. Version 3.1 was used, with the 'GABA+Glx' model applied to the difference spectrum between 2.79 and 4.1 ppm. This model fits the GABA+ peak with a single Gaussian peak around 3.02 ± 0.05 ppm and Glx as a pair of Gaussian peaks at 3.71 ± 0.02 ppm and 3.79 ± 0.02 ppm, and includes terms to characterize the baseline. The Gannet baseline is modelled as a linear slope with sinusoid and cosine terms, periodic at 2.62 ppm (i.e., $A * (f - f_0) + B * \sin( \pi * f / 1.31 / 4 ) + C * \cos( \pi * f / 1.31 / 4 )$ where $f$ is the frequency (in ppm) and $f_0$ the offset to the first modelled Glx peak (±3.71 ppm).

The default Gannet processing pipeline applies zero-filling to 0.062 Hz spectral resolution, and line-broadening of 3 Hz. Both these steps were omitted in the present study.

##### H.1.3 LCModel

LCModel [13] was invoked with the following parameters for modelling GABA-edited difference spectra, to achieve a relatively stiff but non-zero cubic spline baseline model:

- `SPTYPE='mega-press-3'` sets the "special type" appropriately for MEGA-PRESS. This implies a flat baseline (`NOBASE=T`), the use of NAA @ 2.01 ppm as a reference for basis set scaling (`wsmet='NAA', wsppm=2.01`), and default correction factor for attenuation of NMR-visible water, ATTH2O = 0.43 (suitable for TE = 68 ms).

- `NOBASE=F` to re-enable baseline modelling (which is disabled by default for MEGA-PRESS when using `SPTYPE=mega-press-3`),

- `DKNTMN=0.6` to set the baseline knot spacing to 0.6 ppm (the default value is 0.15). This value was varied in the exploratory analysis, Supplementary Section G.
- `PPMST=4.2, PPMEND=0.2` model on the range 0.2-0.4 ppm…
- `PPMGAP(1,1)=1.95, PPMGAP(2,1)=1.2` …but exclude the 1.2-1.95 ppm range from modelling
- `DOWS=T` do water scaling
- `DOECC=F` disable ECC

For the simulated datasets containing only GABA signal, it was additionally necessary to disable frequency referencing: `DOREFS=F,F` and `PPMSHF=0.0`.

##### H.1.4 Osprey

Osprey [14] implements a frequency domain basis set fit; version 1.0.1.1 was used, with default fit and reconstruction settings: separate fit on the range 0.2-4.2 ppm, incorporating macromolecule and lipid components in the basis set but without adding a co-edited MM3 peak. Knot spacing for the spline baseline was set at 0.6 ppm. Although Osprey implements a number of soft constraint models, in this instance the MM3co basis set component was incorporated without additional constraints on amplitude.

##### H.1.5 spant

Spant was configured with automatic baseline flexibility `aic_smoothing_factor`=20, as detailed in Supplementary Section E.

##### H.1.6 Tarquin

A local build of Tarquin [15,16] v4.3.11 was used; processed data and per-vendor basis sets were supplied to Tarquin in the corresponding LCModel file formats.

NAA was used as a reference for basis set scaling. Appropriate TE, sweep width and centre frequency (`echo`, `fs`, `ft`) were specified on the command line. Automatic phasing and referencing (`auto_phase` and `auto_ref`) were disabled. The start point for analysis of the difference spectra (`start_pnt`) was set at 5 ms (calculated as 0.005 times the sample frequency). Eddy-current correction was disabled (`--water_eddy false`). HSVD was performed (per defaults) to remove residual water.

Typical parameters for Tarquin, modelling GABA-edited difference spectra:

```
tarquin
--format lcm
--input S12_GABA_68.diff.RAW
--input_w S12_GABA_68.diff.REF

--echo 0.068
--fs 5000
--ft 127714400
--start_pnt 25
--pul_seq mega_press

--auto_phase false
--auto_ref false
--water_eddy false

--w_att 0.76
--w_conc 35880

--ext_pdf true
--output_txt S12_GABA_68.diff.tarquin.txt
--output_csv S12_GABA_68.diff.tarquin.csv
--output_pdf S12_GABA_68.diff.pdf
--output_fit S12_GABA_68.tarquin.fit
--basis_lcm ge_diff_4096_5000.basis
```

#### H.2 Creatine Reference: Edit-OFF sub-spectrum modelling

For modelling of the edit-OFF sub-spectra (for all algorithms except for Gannet), a standard simulated basis set specific to each hardware vendor was adopted. These were derived for edit-OFF sub-spectra using a similar method to that used for the difference spectra (as detailed in Section H.1), incorporating a default set of basis components as defined within Osprey: ascorbate (Asc), aspartate (Asp), creatine (Cr), a negative correction term for the Creatine $CH_2$ singlet around 3.94 ppm (CrCH2), γ-aminobutyric acid (GABA), glycerophosphocholine (GPC), glutathione (GSH), Glutamine (Gln), Glutamate (Glu), water ($H_2O$), myo-inositol (Ins), lactate (Lac), N-acetyl aspartate (NAA), n-acetylaspartylglutamate (NAAG), phosphorylcholine (PCh), phosphocreatine (PCr), phosphoethanolamine (PE), scyllo-inositol (Scyllo), taurine (Tau), tyrosine (Tyros), and a series of macromolecule and lipid components: $MM_{0.9}$ $MM_{1.2}$ $MM_{1.4}$ $MM_{1.7}$ $MM_{2.0}$ $Lip_{0.9}$ $Lip_{1.3}$ $Lip_{2.0.}$

##### H.2.1 FSL-MRS

Edit-OFF sub-spectra were quantified with the standard edit-OFF basis set, using Cr+PCr as an internal reference for basis set scaling. For expedience, the default Newton optimisation algorithm was used; otherwise, all parameters were the same as for the difference case.

```
--basis             ge_off_8192_4000.basis
--data              S12_GABA_68.off.nii.gz
--h2o               S12_GABA_68.ref.nii.gz
--output            S12_GABA_68.off
--overwrite
--TE                68
--TR                2.0
--tissue_frac       .5 .5 0
--verbose
--report
--combine           NAA NAAG
--combine           Glu Gln GSH
--combine           Cr PCr
--combine           GPC PCh

Defaults:
--internal_ref      Cr PCr
--algo Newton
--ppmlim:           (0.2, 4.2)
--baseline_order:   2
--metab_groups:     0
--h2o_scale:        1.0
```

##### H.2.2 Gannet

In addition to the GABA+Glx model, Gannet independently fits a Lorentzian model to NAA around 2.01 ± 0.04 ppm, and a dual-Lorentzian model for Choline and Creatine, the latter centred at 3.02 ppm with a fixed 0.18 ppm separation, all from the edit-OFF sub-spectrum. A minor local modification was made to additionally yield water-referenced estimates from the existing Cr and NAA fits (usually only reported for those metabolites contained in the difference spectrum).

##### H.2.3 LCModel

Basic configuration was similar to that used for the difference spectra, with two key exceptions. `PPMGAP` was not defined – which implies fitting over the full defined fit range (0.2 - 4.2 ppm), the default mode of operation. `SPTYPE` and `DKNTMN` were also not defined, again implying a default fit: a regular spline baseline was modelled (with default knot spacing 0.15 ppm), Creatine was used for internal basis set scaling, and LCModel's default set of soft constraints were adopted.

##### H.2.4 Osprey

Edit-OFF sub-spectra were modelled with default parameters, in the same invocation of OspreyFit as for the difference spectra.

##### H.2.5 spant

Edit-OFF spectra were quantified using default parameters for spant.

##### H.2.6 Tarquin

Edit-OFF sub-spectra were quantified with HSVD water removal disabled (`--water_width 0`) but all other parameters equivalent to those used for the difference spectra.

```
tarquin

--format      lcm
--input       S12_GABA_68.off.RAW
--input_w     S12_GABA_68.off.REF

--echo        0.068
--fs          5000
--ft          127714400
--auto_phase  false
--auto_ref    false
--water_eddy  false
--water_width 0

--w_att       0.76
--w_conc      35880

--output_txt  S12_GABA_68.off.txt
--output_csv  S12_GABA_68.off.csv
--output_pdf  S12_GABA_68.off.pdf
--output_fit  S12_GABA_68.off.fit
--ext_pdf     true

--basis_lcm   ge_off_4096_5000.basis
```
